# Uranium mining fuels evolution in deep groundwater microbiomes

**DOI:** 10.1101/2025.07.27.666752

**Authors:** Wei Xiu, Till L.V. Bornemann, Tianjing Zhang, André R. Soares, Luping Xie, Julian M. Künkel, Guoxi Lian, Haicheng Weng, Chongsheng Lu, Bing Yang, Ruixuan Gai, Zhipeng Gao, Di Zhang, Yuan Yuan, Zhiming Du, Qingyin Xia, Linxing Chen, Jonathan R Lloyd, Hailiang Dong, Xuebin Su, Huaming Guo, Alexander J. Probst

**Author notes:** to whom correspondence should be addressed: Prof. Alexander J. Probst; Prof. Huaming Guo.

## Abstract

Deep-biosphere microbes have been underpinning global biogeochemical cycles throughout Earth history, yet their evolutionary responses to long-term anthropogenic disturbance remain poorly understood. Neutral-pH in-situ leaching (ISL), the dominant uranium-mining strategy, imposes persistent radiochemical and redox stress offering rare natural experiments across decades for observing subsurface microbial evolution. Here, by exploiting metagenomics and metatranscriptomics in examining microbial responses through various mining stages, we show that neutral U ISL causes substantial changes in microbial communities spanning 2,294 strains accompanied by diversification and selection of active microbial species. These changes occurred during elevated dissolved uranium and radiological activity which correlated with a substantial transcriptional change in energy metabolism, and with an overexpression of specific genes for oxidative-stress defence and DNA-repair pathways. Increased nucleotide diversity in 101 out of 392 species clusters and nonsynonymous/synonymous polymorphism ratios greater than one in 139 species clusters were predominantly observed in metatranscriptomes with elevated radiation, highlighting positive selection of transcriptionally active populations. These findings demonstrate that neutral U ISL drives functional and genetic diversification of subsurface microbiomes, revealing a dynamic and evolutionarily responsive deep biosphere.

## Introduction

Microbial consortia in the deep subsurface, adapted to nutrient-scarce, anoxic environments^1–4^, mediate critical biogeochemical processes and help maintain groundwater quality^5–8^. However, anthropogenic stresses arising from mining, groundwater extraction, and energy production are perturbing these fragile environments at increasing rates^9^. Recent studies have shown that environmental stresses, including heavy metal contamination and altered pH, affect groundwater microbial communities^10,11^. Mining operations are particularly disruptive, as they rapidly subject pristine subsurface ecosystems to heavy metals, radioactive contaminants, altered redox conditions, and electrochemical gradients^9^. As mining encroaches on fragile groundwater ecosystems, resolving the evolutionary responses of subsurface microbiomes is essential for predicting the long-term consequences of resource extraction. Notably, a recent study in the abyssal seafloor showed that deep-sea mining can cause biological disruptions persisting for decades^12^, underscoring the need to assess long-term impacts of subsurface resource extraction on microbiomes. Although distinct from the deep subsurface examined here, this finding underscores how resource extraction can impose long-lasting impacts on microbial communities in Earth’s hidden biomes.

*In*-situ leaching (ISL), a widely used uranium (U) mining technique, is based on injecting oxidizing agents into aquifers to mobilize U by oxidizing insoluble U(IV) to soluble U(VI). In addition to fundamentally altering groundwater chemistry^13^, this activity imposes profound chemical and radiological stresses on microbial communities that have evolved over a long time under stable and reducing conditions. Whereas the environmental consequences of ISL are well documented mostly via 16S rRNA gene analysis^11,14–16^, the evolutionary responses of microbial communities to these stressors remain poorly understood. Indeed, while mining-driven alterations to groundwater chemistry are expected to disrupt the microbial community structure, functional potential, and genetic diversity^17–20^, the question remains as to whether anthropogenic stress can drive microbial evolution in deep aquifers.

To this end, we implement metagenomic and metatranscriptomic strategies to examine microbial community dynamics and evolutionary adaptations at the first large-scale, neutral-pH ISL operation in China, which uses CO_2_ and O_2_ as leaching agents and targets a sandstone-hosted uranium deposit. It involved sampling aquifers spanning pre-mining, active mining, and post-mining phases. Leveraging this rare sampling opportunity to study microbial adaptation as a response to mining and associated geochemical changes, we show that increases in Eh and the accompanying rise in oxidative stress, including U release and associated radioactivity, are linked to restructuring of microbial communities, enhanced oxidative-stress and DNA-repair responses, and alterations in genetic diversity. Elevated pN/pS ratios (the ratio of the rates of non-synonymous (pN) and synonymous (pS) polymorphism) and shifts in nucleotide diversity implicate mining-induced environmental gradients in significantly altering microbial evolutionary processes. Our findings yield new insights into how deep subsurface microbiomes structurally and evolutionally adapt to anthropogenic stress, bearing implications for global biogeochemical cycles, ecosystem resilience, and bioremediation strategies. The findings of this investigation underscore the consequences of uranium ISL on the evolution of subsurface life, offering a novel framework for appraising microbial resilience and adaptation in mining-impacted deep subsurface ecosystems.

## RESULTS

### Geochemical and radiological alterations during and following Uranium ISL

Neutral pH uranium *in-situ* leaching (neutral-pH ISL) commences while the aquifer is in its natural state (pre-mining phase), proceeds through the injection of leaching solutions (mining phase), and concludes with natural restoration efforts upon cessation of mining activities (post-mining phase; *Figs. 1a-b*). We characterized the geochemical and radiological shifts across all phases of neutral-pH ISL by collecting 80 groundwater samples from four distinct mining sites in Qianjiadian sandstone-hosted uranium deposit in China, tapping aquifers at 220-460m depth below surface (*Figs. 1b-c*). These sites underwent neutral-pH ISL for varying durations, creating a unique time-resolved dataset with considerable variation in their redox conditions, from neutral-reducing pre-mining (BG, baseline groundwater) to weakly acid-oxidizing during mining (MT, mining groundwater), and back to neutral-reducing post-mining (PMI, PMIS, and PMS, representing initial, intermediate, and stable post-mining phases, respectively; *Fig. 1c*).

**Figure 1.**
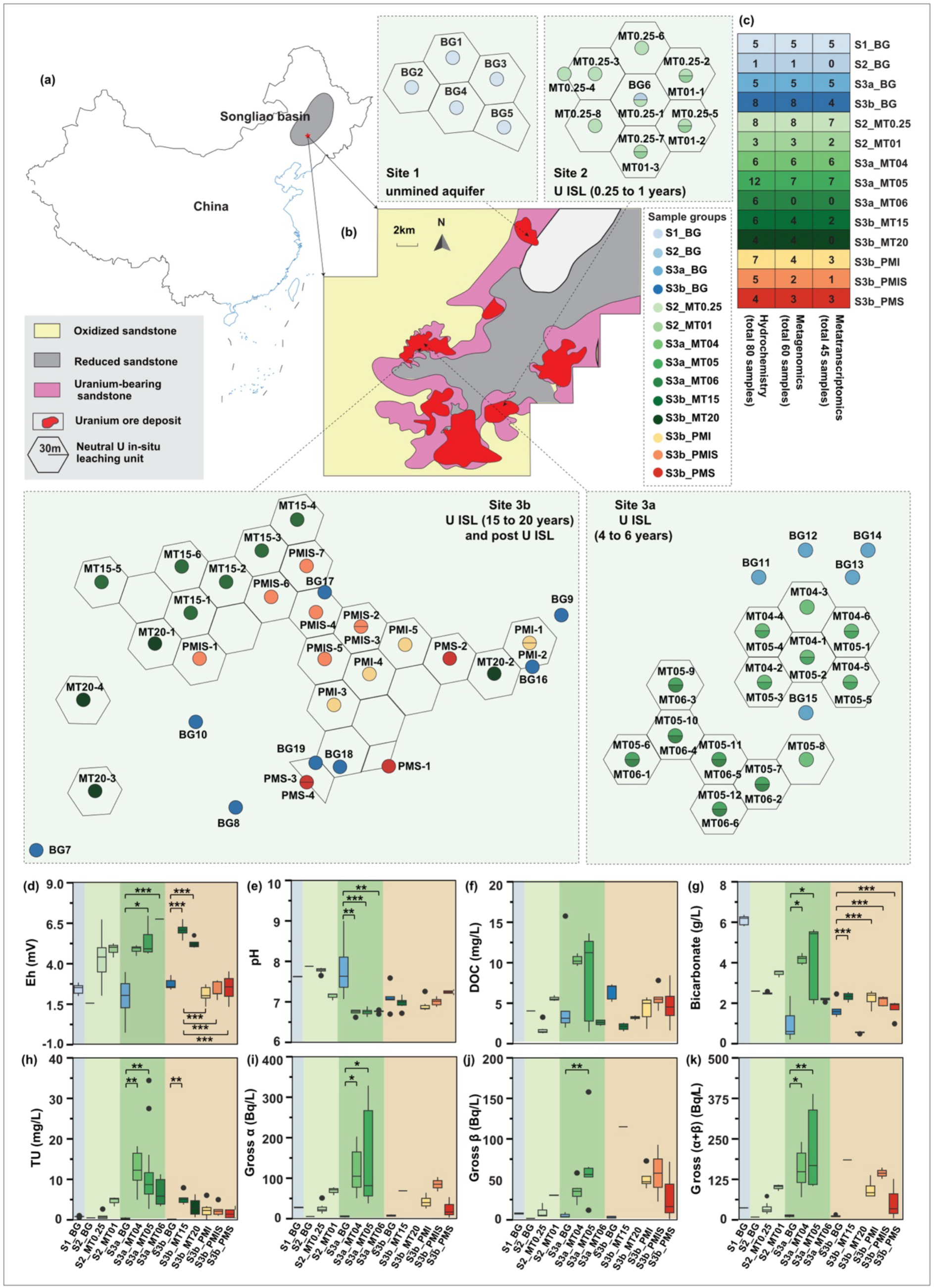
Sampling locations and study overview and geochemical profiles of groundwater across neutral-pH ISL phases. (a) Geographic location of the Songliao Basin and broader regional context, modified from a previously published source^24^. (b) *Schematic layout of study sites along a neutral-pH ISL chronosequence: Site 1 (unmined aquifer, S1_BG); Site 2 (unmined aquifer, S2_BG; recent ISL at 0.25-1 years: S2_MT0.25 and S2_MT01); Site 3a (unmined aquifer, S3a_BG; intermediate ISL, 4-6 years: S3a_MT04, S3a_MT05, and S3a_MT06); and Site 3b (unmined aquifer, S3b_BG; long-term ISL, 15-20 years: S3b_MT15 and S3b_MT20; and decommissioned neutral-pH ISL under iron-reducing, S3b_PMI, mixed iron- and sulfate-reducing, S3b_PMIS, and sulfate-reducing, S3b_PMS conditions). Wells shown with two names indicate that the same well was sampled at different time points. The background map shows oxidized sandstone (yellow), reduced sandstone (grey), uranium-bearing sandstone (pink), uranium ore deposits (red), and neutral-pH ISL units (arrow pointers). (c) Numbers of groundwater samples collected for hydrochemical, metagenomic, and metatranscriptomic analyses across stages of neutral-pH ISL. Boxplots show geochemical parameters through the stage of neutral-pH ISL, including pre-mining (BG), mining (MT), and post-mining (PM). Panels (d – k) represent Eh, pH, DOC, bicarbonate, TU, gross α, gross β, and gross (α+β), respectively. Statistical significance was determined by ANOVA with Tukey’s or Kruskal-Wallis with Dunn’s test (BH-corrected), depending on data normality. Colors indicate study sites: light blue (Site 1), light green (Site 2), medium green (Site 3a), and beige (Site 3b). Blue boxplots within the studied sites (S2_BG, S3a_BG, and S3b_BG) represent the geochemical parameters from unmined aquifer samples*.

Geochemical and radiological profiles shifted markedly across the pre-mining, mining and post-mining stages of neutral-pH ISL. During the pre-mining phase, groundwater was neutral-reducing (Fig. *1d*), with low total dissolved U (TU, 0.44 ± 0.31 mg/L; Fig. 1h). The aquifer was enriched in electron donors, including dissolved organic carbon (DOC, 5.26 ± 3.51 mg/L), Fe(II) (0.85 ± 1.11 mg/L), ammonium (0.73 ± 0.70 mg/L), and sulfide (0.92 ± 1.18 mg/L) (Fig. 1f,S1), conditions that favour the reductive immobilization of U and are consistent with ongoing U mineralization under reducing conditions. In the mining phase, the injection of CO_2_ and O_2_ coincided with a clear oxidative shift, marked by significant increases in Eh (Fig. *1d*) and a rapid rise in TU (Fig. *1h*), which peaked after approximately four years (12.44 ± 5.04 mg/L in S3a_MT04; S3a_MT04, *n* = 6 vs. S3a_BG, *n* = 5; Kruskal–Wallis with Dunn’s post hoc test, *p* = 0.003) before declining gradually from MT5 to MT20 (Fig. *1h*). Radiological activity increased in parallel, with elevated gross β and gross (α+β) signals (Figs. *1i–k*). Bicarbonate concentrations also rose significantly (Fig. *1g*), and both sulfate and nitrate increased in mid-mining stages (Fig. *S1* and Table *S1*). Following neutral-pH ISL decommissioning, groundwater shifted toward reducing conditions, with significant decreases in Eh across MT15 compared with all post-mining groups (S3b_PMI, S3b_PMIS, and S3b_PMS; Kruskal-Wallis with Dunn’s post hoc test, *p* < 0.001; Fig. 1d) and corresponding declines in TU (S3b_MT15 vs. S3b_PMS; ANOVA with Tukey’s test, *p* = 0.036; Fig. *1h*). This transition from oxic to anoxic conditions was further reflected by decreases in sulfate, and sulfate/Cl ratio, whereas Fe(II) concentrations increased, although these changes were not statistically significant (Fig. *S1* and Table *S1*). TU concentrations did not return to pre-mining levels (*Fig. 1h*), indicating incomplete restoration of the aquifers’ original geochemical conditions.

Correlation analysis identified significant Spearman associations among major geochemical and radiological parameters (Fig. *S2*). TU showed strong positive correlations with bicarbonate (*ρ* = 0.43, n = 80, *p* < 0.0001), sulfate (*ρ* = 0.65, n = 80, *p* < 0.0001), and nitrate (*ρ* = 0.69, n = 80, *p* < 0.0001). In addition, TU exhibited positive correlations with gross α (ρ = 0.24, n = 80, *p* < 0.0001), gross β (*ρ* = 0.24, n = 80, *p* < 0.0001), and gross (α+β) (*ρ* = 0.24, n = 80, *p* < 0.0001). These relationships summarize the major covariation patterns among dissolved uranium, radiological activity, and oxidized anions across all samples.

### Metagenome-assembled genomes reveal microbiome changes and enhanced co-occurence resulting from in-situ mining activity

Sixty groundwater samples representative of the pre-mining, mining, and post-mining phases across more than 20 years of neutral-pH ISL were metagenomically sequenced (Figs. *1b,c*). Exceeding 75% of sequencing coverage, based on Nonpareil estimates, indicated a sufficient depth to accurately reflect the aquifer microbiome diversity (Fig. *S3*). To assess both bacterial and archaeal diversity, we focused first on analyzing the ribosomal protein S3 gene (*rpS3*)^21^ yielding 12,972 sequences that clustered (at 99% similarity) into 6,487 representative sequences spanning 215 distinct phyla (21 archaeal and 194 bacterial; Fig. *S4*). Microbial diversity also showed strong positive associations with uranium concentration and radioactivity, with prolonged neutral-pH ISL (S3a_MT04 and S3a_MT05) exhibiting significantly higher Shannon and Invsimpson indices (ANOVA with Tukey’s post hoc test, *p* < 0.05; Fig. *S5*). Groundwater microbial community composition and alpha diversity, based on *rpS3* gene profiles, were strongly influenced by geochemical gradients, particularly radiochemical activity, with higher diversity and distinct community structures observed under elevated stress conditions (*Figs. S5,S6)*, as supported by *Bioenv* analysis (*R*² = 0.051).

We reconstructed 4,075 medium-quality metagenome-assembled genomes (MAGs; completeness ≥ 50%, contamination < 10%)^22^, from which 2,294 strain-cluster representative MAGs (strain-cluster rMAGs, 99% ANI) were obtained, with *Acidobacteriota* representing the most abundant phylum (Figs. *3a-b*). These included 1,239 high-quality and 1,155 medium-quality strain-cluster rMAGs bearing a median completeness of 94.2%, contamination level of 1.3%, and N50 of 47,633 (Figs. *3c-d* and *Table* S2). Consistent with known genome-size differences between domains, archaeal strain-cluster rMAGs had substantially smaller genome sizes than bacterial ones (two-tailed Welch’s t-test, *p* < 0.0001; Fig. *3e*), the largest genome representing an organism from the family *Polyangiaceae* (12.28 Mbp). These strain-cluster rMAGs spanned 63 of the 215 *rpS3-*detected phyla (29.3% of total phylum richness) yet represented 75.1% of the *rpS3*-based phylum-level abundance, thereby providing a robust representation of the predominant microbial communities (*Fig. S7*). The remaining 70.7% of phyla, comprising the 24.9% of *rpS3*-based abundance, were not captured in the strain-cluster rMAGs, potentially reflecting their lower abundance and/or genomes that did not meet medium-quality criteria.

**Figure 3.**
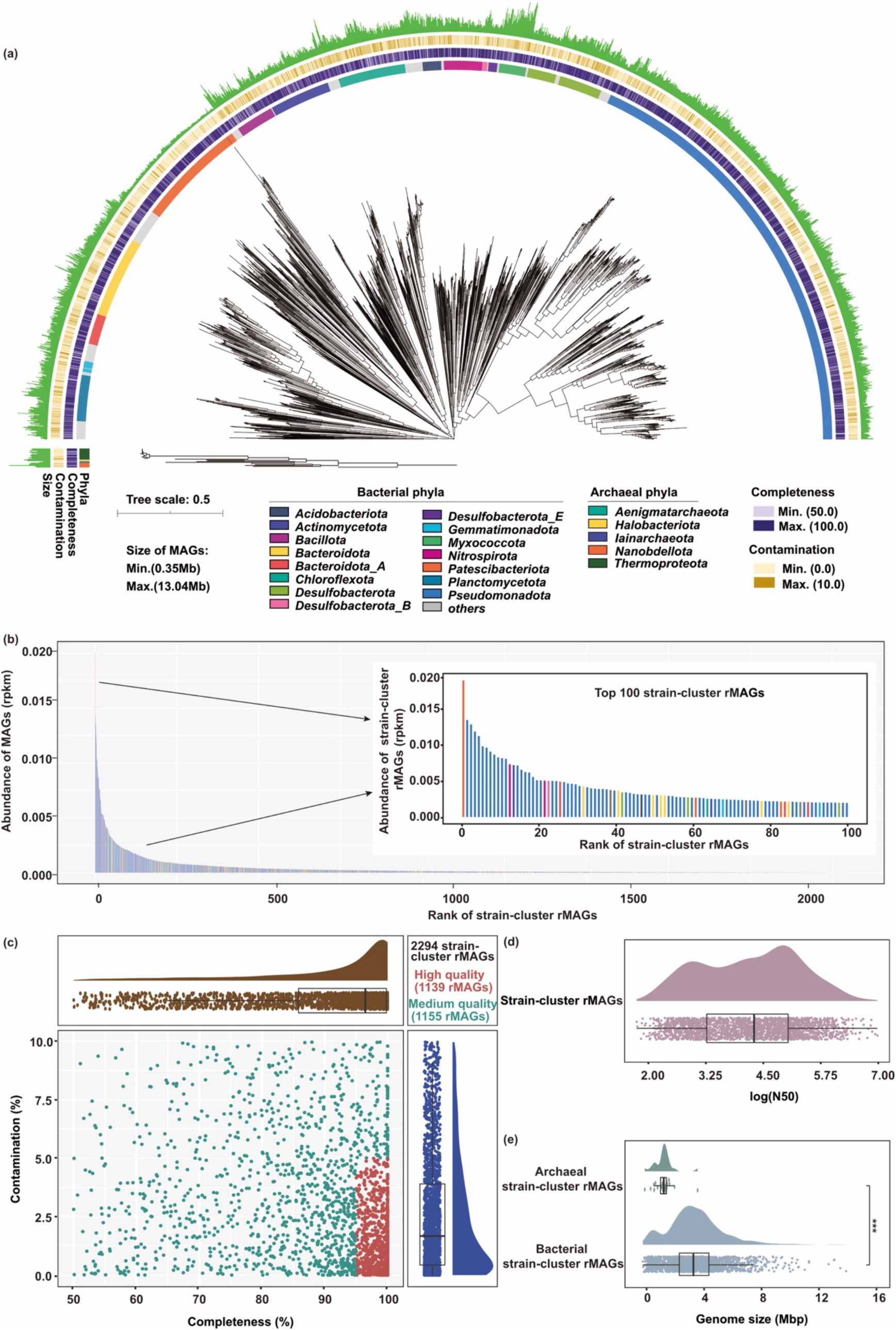
Phylogenetic diversity, abundance patterns, and genomic characteristics of dereplicated strain-cluster representative MAGs. *(a) Maximum-likelihood phylogenetic tree of 2,294 dereplicated bacterial and archaeal strain-cluster rMAGs constructed. Outer rings indicate strain-cluster rMAG taxonomic affiliation at the phylum level (innermost ring), estimated completeness, contamination, and genome size (outermost green bars). (b) Rank abundance of strain-cluster rMAGs across phyla, with the top 100 depicted in more detail. The displayed abundances represent the average strain-cluster rMAG abundances across all samples. (c) Strain-cluster rMAGs completeness and contamination by quality tier. (d) Distribution of log(N50) values of strain-cluster rMAGs. (e) Genome size comparison between archaeal and bacterial strain-cluster rMAGs. Note that two rMAGs showed long-branch or model-sensitive placements; these were retained for completeness but do not affect the overall phylogenetic structure*.

Microbial community composition, based on the relative abundance of strain-cluster rMAGs, shifted markedly along the neutral-pH ISL chronosequence (Fig. *S8*), as evidenced by sample-to-sample Spearman correlation analyses (Fig. *4a*) and non-metric multidimensional scaling (NMDS; Fig. *4b*). Communities from pre-mining, mining, and post-mining stages formed distinct clusters, with analysis of similarities (ANOSIM) indicating significant separation between stages (Fig. *4c*), reflecting divergent community trajectories during and after ISL. Network analysis of relative abundances of strain-cluster rMAGs revealed positive correlations, accounting for 90.39% of all edges, defined as significant correlations between taxa, dominating microbial interactions (Fig. *4d*). Four distinct modules (from Mod1 to Mod4) were identified. Mod1 remained stable across all ISL stages, consistent with a core community. Mod2 and Mod4 increased only during the early ISL stages, reflecting short-term adaptive responses. In contrast, Mod3 showed a sustained increase during the mining and post-mining phases, indicative of recovery processes and stress-driven selection for taxa better adapted to prolonged radiochemical and redox stress (Figs. 4f-i). Within-module degree (Zi) and particitation coefficient (Pi) analysis identified two module hubs (two strain-cluster rMAGs classified within the *Algoriphagus* and *Paracoccus*) that structured interactions within Mod4, and three connector hubs (three strain-cluster rMAGs classified within *Pseudotabrizicola*, *Burkholderiaceae* MESE01, and *Nocardioides*) that linked multiple modules. In addition, 341 peripheral connectors (Zi ≤ 2.5 and Pi > 0.62) were dominated by Mod3 (175 strain-cluster rMAGs) and Mod2 (129 strain-cluster rMAGs) (Fig. *4e*), indicating that most cross-module interactions are mediated by diffuse peripheral taxa rather than by central hub nodes. These results highlight the significant shifts in microbial community structure, diversity, and ecological interactions imposed by neutral-pH ISL. The observed tendency towards positive interactions emphasizes the dynamic and evolving nature of subsurface microbiomes in response to acute environmental stress.

**Figure 4.**
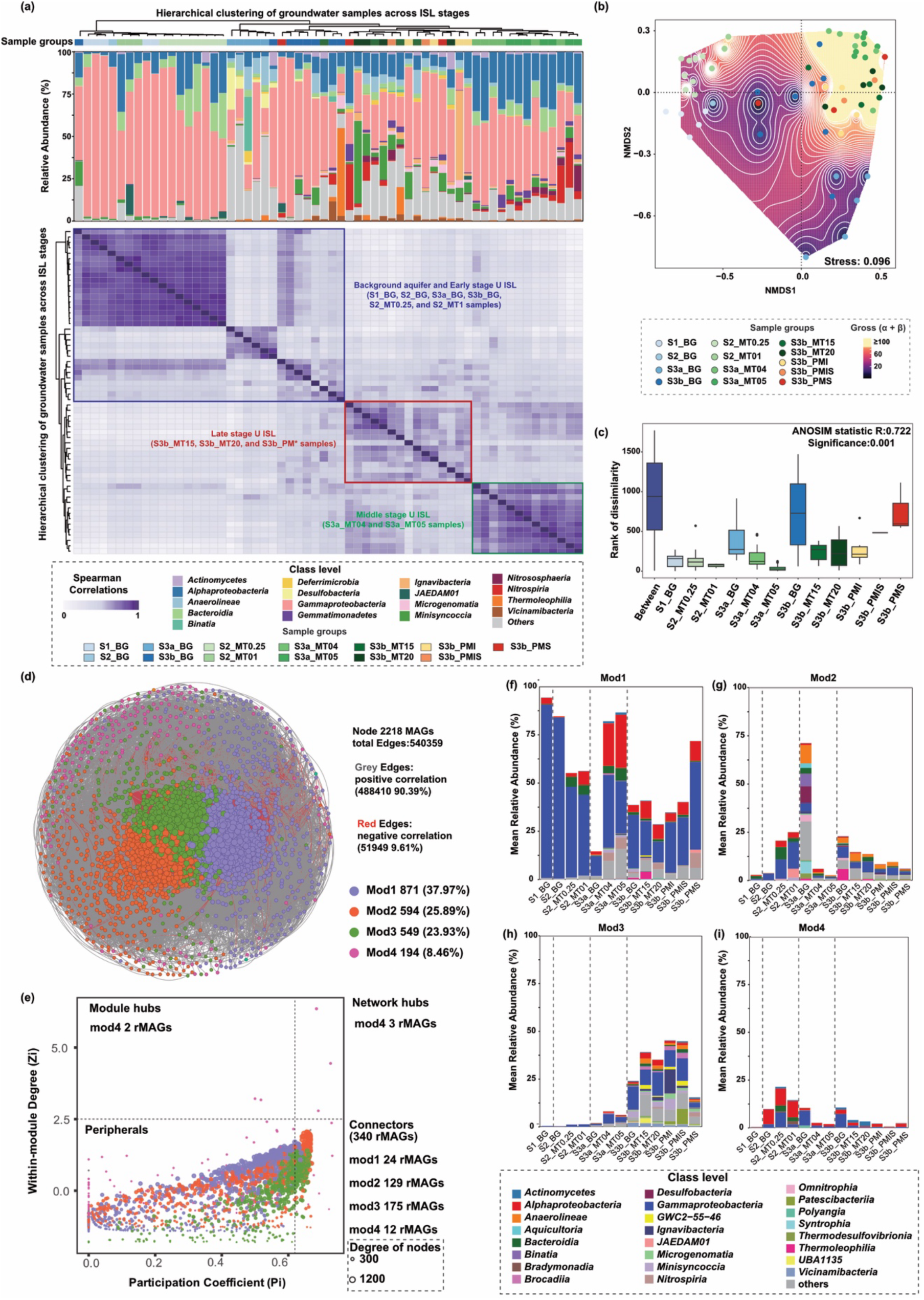
Microbial community structure and modularity across neutral-pH ISL phases. (a) Heatmap of sample-to-sample Spearman’s rank correlations of strain-cluster rMAG abundances across early (blue), mid (green), and late (red) neutral-pH U ISL stages. The accompanying bar chart shows the top 12 classes across samples, *arranged in the same sample order as in the heatmap. (b) Non-metric multidimensional scaling (NMDS) plot of community structure based on strain-cluster rMAGs (stress = 0.096). Background colors represent gross (α+β) radiation levels across all groups (interpolated across the ordination space using inverse distance weighting (IDW). Gross (α+β) radiation levels below 100 Bq/L are shown as a continuous gradient, while gross (α+β) radiation levels ≥100 Bq/L are displayed as a solid yellow. White contour lines and labels indicate gross (α+β) isopleths (Bq/L). (c) Analysis of Similarity (ANOSIM) showing significant differences between groups. (d) Network of strain-cluster rMAGs co-occurrence, with positive (grey) and negative (red) Spearman correlations grouped into four modules. Node size is proportional to the degree (number of connections). (e) Within-module degree (Zi) and participation coefficient (Pi) analysis of the network to identify the peripherals, connectors, module hub and network hubs. The size and color of the nodes are proportional to its corresponding degree and module in the network, respectively. (f-i) Distribution of the abundance of module-specific taxa across neutral-pH ISL phases for Mod1 to Mod4*.

### Neutral-pH ISL impacts gene expression linked to DNA repair, oxidative stress, and biogeochemical cycling

To resolve microbial functional responses to prolonged neutral-pH ISL, we analyzed metatranscriptomes from 45 genomically resolved communities spanning the pre-mining, mining, and post-mining phases. Integration of transcript profiles with 2,294 functionally annotated strain-cluster rMAGs identified 350,588 of 8,422,368 representative genes showing at least two-fold upregulation relative to the mean expression of 30S ribosomal protein genes, which served as an internal benchmark for transcriptional activity. The transcriptional landscape of entire communities based on strain-cluster rMAGs showed clear patterns of transcriptionally up-regulated populations that are specific for the sample groups across the mining sites (Fig. *S9*). Among 350,588 upregulated genes of the rMAGs, 5,677 genes were associated with carbon, hydrogen, oxygen, nitrogen, sulfur, and iron biogeochemical transformations, 4,624 with DNA repair pathways, and 2,760 with oxidative stress responses (Figs *S10-S17*), highlighting broad activation of redox and genome-stability pathways under ISL-induced chemical and radiological stress.

Regarding microbial activity of biogeochemical transformations, the expression patterns across the 5,677 related genes were strongly structured by geochemical conditions, as revealed by *Bioenv* analysis, which identified pH, Eh, sulfide, sulfate and bicarbonate as the principal geochemical variables shaping the transcriptional structure of overexpressed C-H-O-N-S-Fe metabolic genes across the metatranscriptomes (*R²* = 0.39), with PCoA accounting for 36.9% of the variation among overexpressed genes (Figs. *5a* and *S9*). This pattern is consistent with CO₂ and O₂ injection and with oxidation of ammonium and pyrite as dominant ISL reactions fueling microbial redox metabolism. TU concentrations were moderately and positively correlated with the expression of 20 energy-conserving genes distributed across 220 strain-cluster rMAGs, including *ccoN* (cytochrome *c* oxidase, aerobic respiration; *ρ* = 0.44, *p* = 0.029; encoded in 77 strain-cluster rMAGs), *pmoC* (particulate methane monooxygenase, methane oxidation; 21 strain-cluster rMAGs), *nxrAB* (nitrite oxidoreductase, nitrite oxidation; *ρ* > 0.53, *p* < 0.006; *nxrA* in 21 and *nxrB* in 27 strain-cluster rMAGs), and *soxAYZ* (sulfur oxidation complex, sulfur oxidation; ρ > 0.41, *p* < 0.038; *soxA* in 43, *soxY* in 69, and *soxZ* in 74 strain-cluster rMAGs) (Figs. *5d-f* and *S16-S17*). These relationships highlight a broad activation of energy-yielding redox pathways along radiochemical gradients, underscoring the metabolic coupling of carbon, nitrogen, and sulfur cycling with oxidative stress.

**Figure 5.**
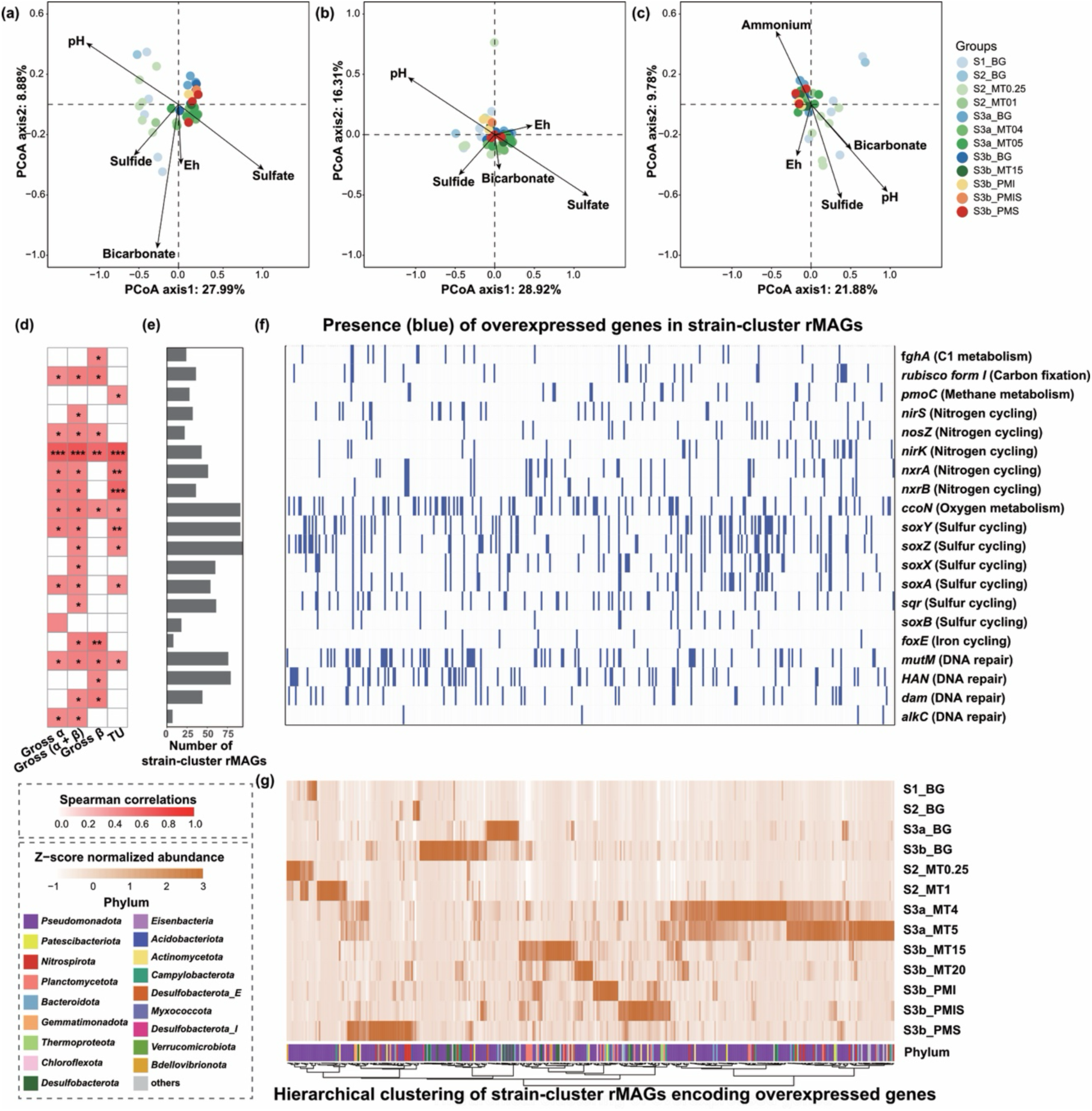
Correlation and environmental drivers of overexpressed genes in metatranscriptomic datasets. (a–c) Principal coordinate analyses (PCoA) based on expression of (a) genes involved in carbon, hydrogen, oxygen, nitrogen, sulfur, and iron metabolism, (b) DNA-repair genes, and (c) oxidative-stress-response genes. Arrows indicate environmental variables significantly correlated with gene-expression patterns, as identified by Bioenv analysis. Points are colored and shaped by sample *groups. (d) Spearman correlations of overexpressed genes positively associated with total dissolved uranium (TU), gross α, gross β, and gross (α+β). (e) Number of strain-cluster rMAGs encoding the corresponding overexpressed genes that are positively correlated with total dissolved uranium (TU), gross α, gross β, and gross (α+β). (f) Presence-absence heatmap of overexpressed genes across strain-cluster rMAGs. (g) Z-score–normalized relative abundances of strain-cluster rMAGs encoding genes positively correlated with TU and radiological activity (gross α, gross β, and gross (α+β)) across neutral-pH ISL stages. rMAGs are clustered by Z-score profiles, with phylum-level taxonomy indicated below*.

In addition to metabolic adjustments, microbial populations showed dynamic transcriptional regulation of DNA-repair and oxidative-stress pathways in response to radiochemical gradients. The *Bioenv* analysis identified pH, Eh, sulfide, ammonium, and bicarbonate as the main variables structuring DNA-repair gene expression (4,624 genes, *R²* = 0.40; PCoA explained 31.7% of variance), whereas pH, Eh, sulfide, sulfate, and bicarbonate were key for the 2,760 oxidative-stress-response genes (*R²* = 0.37; PCoA explained 45.2% of variance; Figs. *5b,c*). At the individual gene level, expression of foxE (c-type cytochrome involved in Fe(II) oxidation) correlated positively with gross β and gross (α+β) (Spearman, *p* < 0.05; 8 strain-cluster rMAGs; Figs. *5b-e*), consistent with Fe redox processes being stimulated by oxygen ingress and the resulting shifts in uranium speciation and radiological activity. Consistent with these environmental associations, expressions of DNA-damage-repair genes, including *mutM* (DNA glycosylase; base-excision repair of oxidized bases; *ρ* = 0.43-0.49, *p* = 0.021-0.047; 68 rMAGs), *dam* (DNA adenine methylase; mismatch-repair guidance; ρ = 0.45-0.51, *p* = 0.016-0.038; 37 rMAGs), and *alkC* (alkylation-repair protein; ρ = 0.45-0.46, *p* = 0.034-0.038; 4 rMAGs), were positively correlated with TU and gross α, β, and (α+β) (Spearman correlations, *p* < 0.05; *Figs. 5b-e*). Notably, *mutM* expression correlated moderately with both TU (ρ = 0.41, n = 26, *p* = 0.0435) and gross (α + β) (ρ = 0.51, n = 2,632, *p* = 0.0084; Fig. *5e*). These coordinated transcriptional patterns indicate widespread modulation of genome-stability and redox-defense mechanisms, coupling Fe redox metabolism, oxidative stress responses, and DNA repair under sustained radiochemical and geochemical stress in the aquifer.

### Uranium ISL-induced stress drives genetic diversification in subsurface aquifers

Within-species selection pressure and genetic diversity were quantified across 2402 members of 392 dereplicated species-cluster rMAGs (95% ANI). Each rMAG was represented by more than three members with >5× genome coverage in 60 metagenomic and 45 metatranscriptomic datasets, spanning neutral-pH ISL phases, enabled direct comparisons of genomic and transcriptional diversity across mining stages.

NMDS analysis of pN/pS ratios revealed a clear progression from background to mining and post-mining samples in both metagenomes (Fig. *6h*) and metatranscriptomes (Fig. *6j*), closely aligned with increasing TU concentrations. Consistent with this ordination, site-level analyses showed that the median metagenomic pN/pS ratios increased in early mining, late mining, and post-mining sites and metatranscriptomic pN/pS ratios exhibited similar but stronger increases (*p* < 0.05, *Figs. 6a-c*). *Bioenv* analysis identified gross α, gross (α+β), Eh, TFe, TU, ammonium, Fe(II), nitrite, and nitrate as the main correlates of metagenomic pN/pS (*R²* = 0.28), and gross β, gross (α+β), TDS, TU, and ammonium for the metatranscriptomes (*R²* = 0.23), indicating that mutation–selection imbalance reflects the combined influence of the oxidizing geochemical regime and co-varying metal and radionuclide stress. Instances of positive selection (pN/pS > 1) were more frequent in the metatranscriptomes (139 MAG clusters) than in the metagenomes (112 MAG clusters) or in both datasets combined (49 MAG clusters; Fig. *S19*), highlighting intensified adaptive pressures on actively expressed genomes.

**Figure 6.**
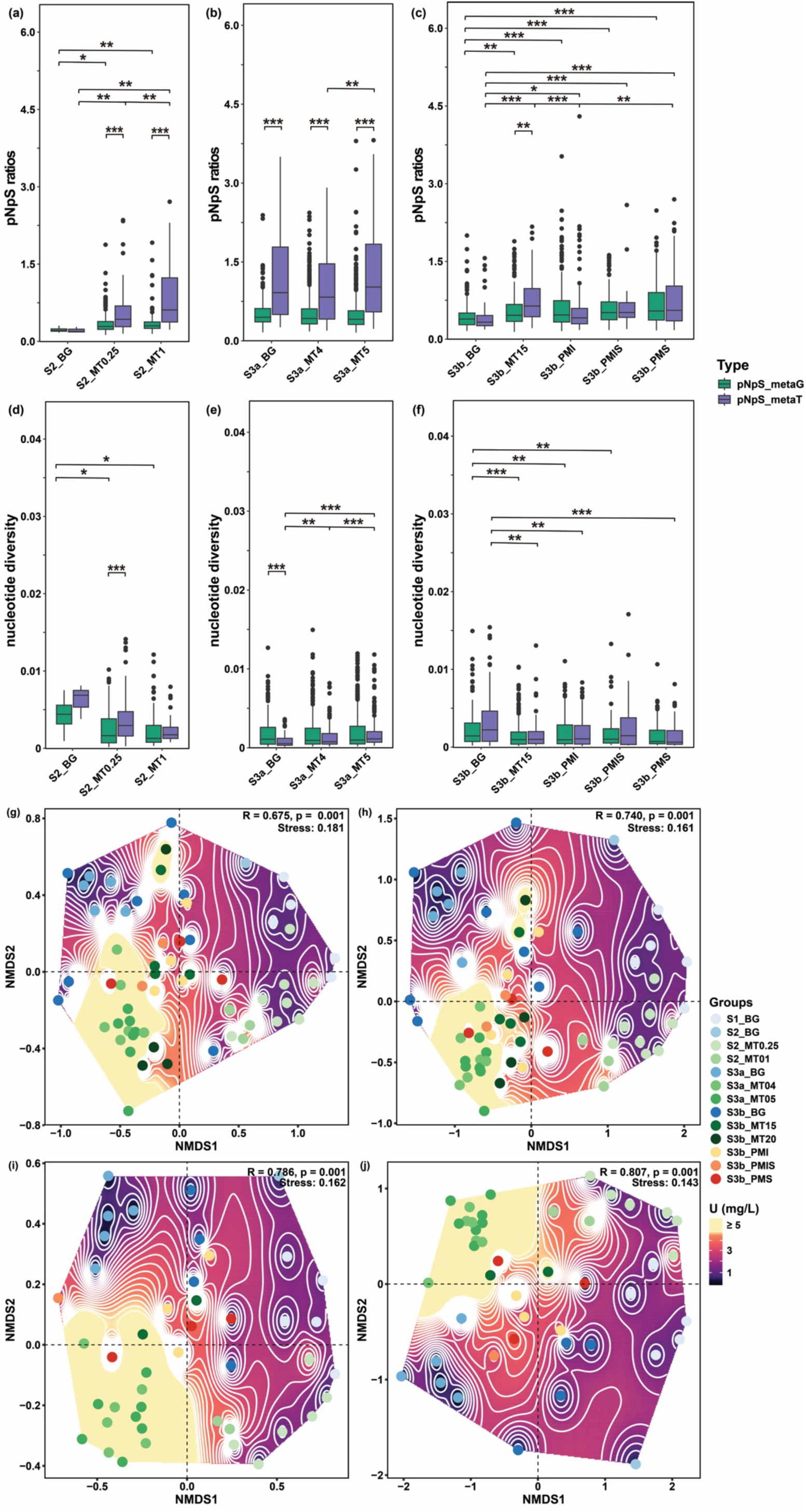
Genome-wide pN/pS ratio and nucleotide diversity across mining stages in metagenomes and metatranscriptomes. (a-c) pN/pS ratio from metagenomic (metaG) and metatranscriptomic (metaT) datasets spanning pre-mining, mining, and post-mining phases; (d-f) Nucleotide diversity from metaG and metaT datasets across neutral-pH ISL stages; (g-j) Non-metric multidimensional scaling (NMDS) plots based on Bray–Curtis dissimilarity showing the relationships between TU and pN/pS ratios or nucleotide diversity derived from metaG (g, h) and metaT (i, j) datasets.Statistical significance was assessed using the Kruskal–Wallis test followed by Dunn’s post-hoc test with Benjamini–Hochberg (BH) correction for multiple comparisons (* p < 0.05, ** p < 0.01, and *** p < 0.001).

To explore whether these shifts in selection were supported by changes in standing genetic variation, we next examined genome-wide nucleotide diversity across species-cluster rMAGs. NMDS ordinations revealed a consistent progression from background to mining and post-mining samples in both metagenomes and metatranscriptomes, closely tracking TU gradients (*Figs. 6g-j*). Metagenomic nucleotide diversity remained relatively stable during the mid-mining phase but declined in early-, late-, and post-mining stages despite sustained TU concentrations. (Figs. *6d-f*). By contrast, metatranscriptomic nucleotide diversity rose sharply during the mid-mining phase at site S3a (*p* < 0.001; Fig. *6e*), coinciding with peak uranium and radioactivity. Bioenv analysis identified radioactivity (gross α, gross β, and gross (α+β)) and geochemical parameters (Eh, TFe, TU, ammonium, Fe(II), and nitrate) as key correlates of metagenomic genome-wide nucleotide diversity (*R²* = 0.25), with comparable associations in metatranscriptomes (gross β, gross (α+β), TDS, TU, and ammonium; *R²* = 0.21). Together, these patterns indicate that prolonged uranium-mining stress maintains elevated genetic variation in transcriptionally active populations, consistent with bet-hedging-like strategies that enhance adaptive flexibility in fluctuating radiochemical environments.

Sixty-five out of 392 species clusters exhibited recurrent increases in nucleotide diversity from background to mining-impacted conditions, including 20 detected in both metagenomic and metatranscriptomic datasets (Figs. *7a,b*). Four representative taxa, *Kuenenia* (rMAG-1), *Gallionella* (rMAG-2 and rMAG-4), and *Parvibaculum* (rMAG-3), persisted across all mining phases and exhibited elevated genome-wide diversity in both metagenomic and metatranscriptomic datasets. Gene-level diversification was dispersed across loci rather than confined to specific genomic regions, with hundreds of variable genes per genome (rMAG-1: 269/1,126; rMAG-3: 95/148; rMAG-4: 213/280; rMAG-2: 295/350) and substantial overlap between genomic and transcriptomic variation, indicating that many variable loci were transcriptionally active, particularly within core functional networks linked to uranium-induced stress responses, including information processing, regulation, and stress adaptation (*Fig. 7c*). *Kuenenia* rMAG-1 exhibited genomic diversification in hydroxylamine dehydrogenase (*hao*) and hydrazine dehydrogenase (*hdh*) genes mediating ammonium oxidation, together with *lexA* (SOS-response regulator involved in DNA damage repair), *old-family endonuclease* (genome maintenance), *lon* (ATP-dependent protease for protein quality control), and *tolC* (outer-membrane efflux channel for metal detoxification). The two *Gallionella* rMAGs displayed contrasting transcriptional responses. The rMAG-2 showed variability in redox-related genes, including cytochrome-c oxidase cbb3-type subunit II (*fixO*), involved in microaerobic respiration and electron transport, and *thioredoxin-dependent peroxiredoxin*, linked to reactive oxygen species metabolism under oxidative stress. By contrast, rMAG-4 exhibited transcript-level variability in outer-membrane porins (*ompC*/*ompF*/*phoE*) associated with metal-regulated permeability, the conjugal transfer protein *traF* involved in horizontal gene transfer, and glycosyltransferases contributing to cell-envelope remodeling. *Parvibaculum* rMAG-3 encoded a GIY–YIG endonuclease, associated with the cleavage of damaged or mismatched DNA, and a Zn-dependent protease involved in protein quality control under oxidative stress, including conditions where metal exposure can exacerbate ROS damage. Together, these diversification patterns suggest that persistent taxa employ distinct, lineage-specific strategies-spanning redox flexibility, oxidative defense, membrane adaptation, and proteogenomic repair, to maintain resilience under the combined chemical and radiochemical stresses imposed by U mining.

**Figure 7.**
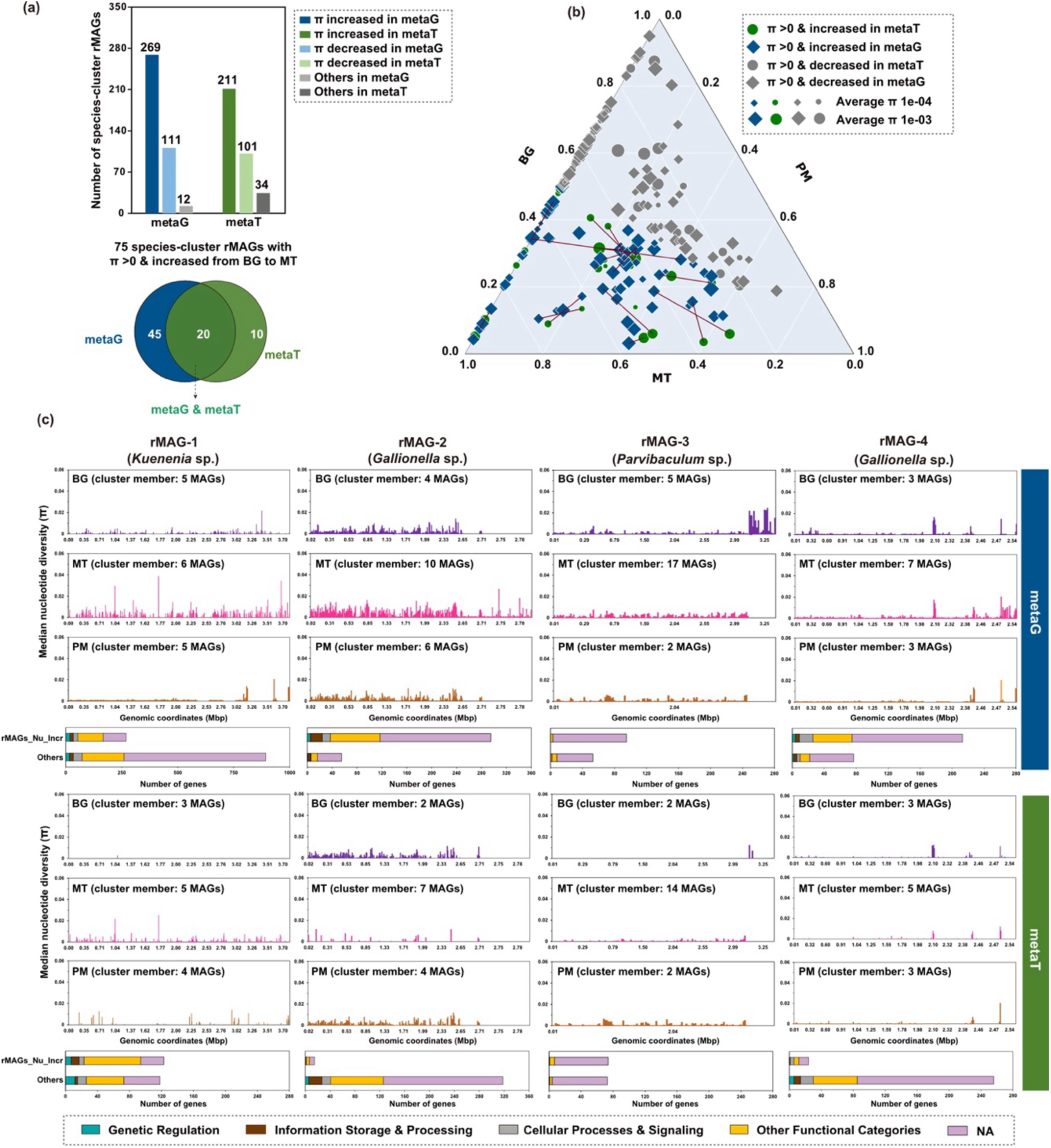
Genome-wide nucleotide diversity patterns across mining stages in representative rMAGs. (a) Bar plots show the total numbers of species-cluster rMAGs with increased or decreased nucleotide diversity (π) from background (BG) to mining (MT) conditions in metagenomic (metaG) and metatranscriptomic (metaT) datasets. Increased π indicates rMAGs with callable nucleotide diversity (≥ 5× coverage) in MT samples but either non-callable diversity (< 5× coverage) or lower π values in BG samples; decreased π follows the inverse criterion. The accompanying Venn plot displays the overlap of rMAGs with increased π (π > 0) between the metaG and metaT datasets. (b) Ternary plot showing the distribution of rMAG-level nucleotide diversity across BG, MT, and post-mining (PM) samples, with points colored by dataset and sized by diversity magnitude. (c) Gene-level nucleotide diversity profiles for four representative species-cluster rMAGs (Parvibaculum, Kuenenia, and Gallionella) across BG, MT, and PM phases. Each profile represents the median π of all cluster members for a given gene, highlighting elevated and widespread diversification during mining (MT), particularly in transcriptionally active genes. Bar plots below each panel show the functional distribution of genes with increased diversity categorized by major gene functions.

## Discussion

The global resurgence of nuclear energy has renewed interest in U ISL, a minimally invasive yet chemically intensive extraction method, poised to dominate future uranium supply chains^23,24^. During mining, strong oxidants drove the oxidation of U(IV) to U(VI) and the formation of soluble uranyl–carbonate complexes, while concurrent oxidation of pyrite and ammonium produced the elevated sulfate and nitrate characteristic of mining phases^25,26^. Following ISL cessation, DOM degradation re-established reducing conditions, accompanied by rising Fe(II) and declines in Eh, TU and sulfate, patterns consistent with Fe(III) and sulfate reduction and the partial reductive attenuation of U(VI)^27^. While such geochemical perturbations and ecological restructuring during U ISL are increasingly recognized^13,17–20,25,27–30^, its evolutionary consequences for subsurface microbiomes remain underexplored. Here, we present metagenomic and metatranscriptomic evidence that neutral-pH U ISL exerts sustained selective pressure on microbial populations, driving *in situ* evolution within deep aquifers.

Neutral-pH ISL establishes persistent redox disequilibria through oxidant injection, mobilizing U, radionuclides, and heavy metals^13,25,27–30^. By examining a neutral-pH ISL area with aquifers having undergone different time periods of leaching (up to 20-years), we reveal that these sustained geochemical disturbances go beyond restructuring of microbial community composition and alter transcriptional activity, along with an increased genetic diversity, illuminating long-term ecological and evolutionary consequences for deep biosphere microbiomes. These transformations are shaped by sustained selection pressures imposed by neutral-pH ISL, including oxidative stress, redox-active metal toxicity, and low-dose radiation, stressors known to destabilize cellular systems and promote adaptive traits linked to stress tolerance, DNA repair, and redox regulation^31–33^. Comparable patterns of stress-driven adaptation have been reported in acid mine drainage biofilms^34,35^, heavy-metal tailings^36,37^, permafrost thaw soils^38^, and deep-sea sediments^39,40^. Consistent with these analogs, we detected increased taxonomic diversity in neutral-pH ISL-impacted aquifers, signatures consistent with stress-induced diversification to increase resilience at the community level^11,41–45^. Temporal succession revealed dynamic microbial responses, with oxidative-stress-tolerant taxa dominating early neutral-pH ISL stages and anaerobic fermenters and metal reducers emerging post-mining, reflecting adaptation to evolving redox conditions.

Crucially, these ecological transitions were accompanied by molecular signatures of adaptation. Genes associated with oxidative damage, heavy metal toxicity, and radiation exposure were broadly overexpressed across all neutral-pH ISL stages (*Figs. S9-S17*), indicating a constitutive stress response. Among 75 species-cluster rMAGs with measurable nucleotide diversity that increased from background to mining phases (Fig. 7a), many exhibited pronounced diversification along the ISL gradient, suggesting that repair mechanisms were insufficient to counteract accumulated genomic damage. Consequently, elevated nucleotide diversity, particularly in mid-mining metatranscriptomes, supports a mutation–repair imbalance in which mutational input exceeded repair capacity, consistent with a previously introduced model of stress-induced mutagenesis^37,46–48^. This model posits that environmentally triggered DNA damage, coupled with error-prone repair, contributes to increased mutation rates. While DNA repair pathways buffer deleterious mutations, they may also limit adaptive potential by constraining the fixation of beneficial variants^49,50^. In our study, concurrent increases in nucleotide diversity and transcriptional upregulation of DNA repair and oxidative stress-response genes indicate that adaptive evolution is concentrated in metabolically active, redox-cycling taxa rather than uniformly across the whole community. This resolves the modest community-wide transcriptional changes yet strong gene-level positive selection signals, highlighting that long-term ISL creates evolutionary hotspots where radiochemical stress and altered substrate availability jointly promote adaptation in key biogeochemical taxa.

Further support for this mechanism comes from elevated pN/pS ratios (>1), a canonical signal of positive selection, observed predominantly in metatranscriptomes (139 species-cluster rMAGs, including 251 and 40 cluster members were oberved in mining and post-mining samples) and only partially overlapping with metagenomic datasets (49 species-cluster rMAGs; Fig. S19). The rMAGs exhibiting elevated nucleotide diversity encompassed phylogenetically diverse taxa and displayed extensive gene-level variation in information processing and regulatory functions, underscoring that selection acts primarily on transcriptionally active populations. This layered response, genome-wide mutation accumulation coupled with transcription-linked selection, highlights the dual role of environmental stress in both generating genetic variability and filtering for functional adaptation, consistent with eco-evolutionary rescue models^51–53^, wherein microbial populations rebound from stress-induced bottlenecks through the acquisition and retention of adaptive mutations.

Collectively, our study provides rare empirical evidence that stress-induced mutagenesis and transcription-linked selection operate within natural deep subsurface microbiomes over decadal timescales. Although such mechanisms are well characterized in laboratory systems^54–60^ and surface environments^46,61–64^, their manifestation in oligotrophic, energy-limited subsurface ecosystems^39,65–67^ has remained largely speculative due to assumptions of metabolic quiescence and evolutionary stasis^40,68–70^. Our findings challenge this paradigm, demonstrating that chronic anthropogenic stress in engineered aquifers can generate sustained mutational input and facilitate directional selection. The convergence of elevated nucleotide diversity, DNA repair activation, and positive selection in expressed genes reveals a coordinated evolutionary response to long-term environmental disturbance. These observations extend core theoretical frameworks, including stress-induced mutagenesis and eco-evolutionary rescue, from controlled systems to complex, anthropogenically modified subsurface environments, redefining the deep biosphere as a dynamic, evolutionarily responsive system.

## MATERIALS AND METHODS

### Study area, sampling strategy, and sample collection

The study area located in the Qianjiadian U ore deposit within the Qianjiadian (QJD) sag of the Kailu Depression in the southwestern Songliao Basin, northeast China, represents a typical sandstone-hosted U deposit^71^. The stratigraphy comprises the Qingshankou (K_2_qn), Yaojia (K_2_y), and Nenjiang (K_2_n) formations, overlain by Neoproterozoic and Quaternary alluvial deposits. The U mineralization is primarily hosted in the K_2_y Formation, which consists of red, yellow, gray, and mineralized gray sandstones, with widespread carbonate cementation (dolomite, siderite, ankerite, and calcite). Pitchblende and coffinite are the dominant uranium minerals, while Fe-Ti oxides, natural organic matter, clay minerals, and pyrite serve as additional U hosts^27,72,73^. Groundwater flow follows a southwest-to-northeast trajectory within a confined Cretaceous sandstone aquifer (approximately 298 m depth below surface, 35 m thick), with permeability averaging 0.145 m/day^25^. The aquifer is hydraulically isolated by a low-permeability mudstone-dominated aquitard both above and below, with limited connectivity to adjacent hydrogeological units.

The Qianjiadian U deposit has been a key site for advancing in-situ leaching (ISL) technology in China, serving as the first large-scale demonstration of CO₂ + O₂ leaching in a deep, low-permeability sandstone-hosted system. During the neutral-pH ISL process, the mixed reagent of dissolved O_2_ (oxidant) and CO_2_ (complexing agent) was injected into the ore-bearing aquifer, and subsequently, U-containing groundwater was pumped out and U was retrieved using ion exchange resins^25,74^. The effluent from the reverse osmosis system was reused as a leaching solution after being mixed with CO_2_ and O_2_ to maintain carbonate buffering and oxidizing conditions^75^. The orebody, located within confined sandstone aquifers at depths ranging from 220 to 460 m, presents complex hydrogeological conditions that influence solute transport, aquifer reactivity, and microbial community dynamics. The unique geological and hydrogeochemical setting of the Qianjiadian deposit, combined with its prolonged history of neutral-pH ISL operations, establishes it as a natural laboratory for investigating subsurface fluid-rock-microbe interactions, U mobilization processes, and the long-term geochemical and microbial responses to neutral-pH ISL as well as post-mining restoration.

To investigate the hydrogeochemical and microbial responses to neutral-pH ISL mining, groundwater samples were collected from the Qianjiadian (QJD) sandstone-hosted U deposit during two field campaigns in October-November 2021 and August-October 2022 (Figure 1). A total of 80 groundwater samples were obtained from sites representing different stages of neutral-pH ISL, spanning ore-bearing aquifers at depths of 220-460 m. Sampling locations encompassed neutral-pH ISL sites spanning multiple operational durations: 0.25 years (S2_MT0.25, 8 samples), 1 year (S2_MT01, 3 samples), 4 years (S3a_MT04, 6 samples), 5 years (S3a_MT05, 12 samples), 6 years (S3a_MT06, 6 samples), 15 years (S3b_MT15, 6 samples), and 20 years (S3b_MT20, 4 samples). Post-mining phases were sampled under distinct redox conditions, including iron-reducing (S3b_PMI, 7 samples), mixed iron- and sulfate-reducing (S3b_PMIS, 5 samples), and sulfate-reducing (S3b_PMS, 4 samples) conditions. Nineteen groundwater samples from unmined aquifer were collected across the chronosequence, including non-mining areas (S1_BG, 5 samples), early-stage mining sites (S2_BG, 1 sample), mid-stage mining sites (S3a_BG, 5 samples), and late-stage/post-mining sites (S3b_BG, 8 samples). Background wells were present at all sites and sampled concurrently with mining and post-mining wells, providing site-specific baseline controls. Detailed hydrogeochemical data for S3b_MT20, S3a_MT04, S3a_MT05, S3b_PMI, S3b_PMIS, S3b_PMS, and S3b_BG are reported in our previous studies^25–28^.

Prior to sample collection using a deep groundwater sampler (Geosub 2, Geotech, USA), monitoring and post-mining wells were pumped for at least 20 minutes until water temperature, electrical conductivity (EC), and oxidation-reduction potential (ORP) stabilized. Groundwater was then pumped after well flushing and approximately 45 ± 0.5 L of water was filtered through 0.2-μm sterile cellulose nitrate membrane filters (Sartorius, Göttingen, Germany) to capture microbial organisms from each groundwater well. All filtered membranes were frozen at -80 °C for high-throughput sequencing. Groundwater samples for physicochemical analysis were collected in 5 L sterile PET bottles and stored at -20 °C.

### Chemical Analysis

Major anions (Cl⁻, NO₃⁻, and SO₄²⁻) were measured using ion chromatography (Dionex ICS-600, Thermo Fisher Scientific, Massachusetts, USA), while major cations (Na⁺, K⁺, Mg²⁺, and Ca²⁺) were determined using inductively coupled plasma optical emission spectrometry (ICP-OES, Optima 8300, PerkinElmer, Massachusetts, USA). Trace elements, including U, Fe, and other transition metals, were quantified using inductively coupled plasma mass spectrometry (ICP-MS, 7900 ICP-MS, Agilent Technologies, California, USA). Alkalinity was measured by Gran titration using a Metrohm 888 Titrando (Metrohm, Herisau, Switzerland). The total organic carbon (TOC) and dissolved organic carbon (DOC) concentrations were determined using a TOC analyzer (TOC-L, Shimadzu, Kyoto, Japan). The gross alpha and gross beta radioactivity of groundwater samples were measured using a low-background α/β meter (LB6008, Beijing Yida Measurement Technology Co., Ltd., Beijing, China) following the thick source method for radioactivity analysis, in which evaporated water residues are deposited onto a planchet to form a uniform thick source for α/β counting. Measurements of gross alpha and gross beta radioactivity were conducted at the China Nuclear Mining Science and Technology Corporation, Beijing, China, to assess radiological characteristics associated with uranium mobilization.

### DNA extraction and sequencing

Total metagenomic DNA was extracted from biomass retained on 0.22-μm sterile cellulose nitrate membrane from 60 groundwater samples, selected from a total of 80 samples retrieved from different neutral-pH ISL stages in ore-bearing aquifers (see above and Figure *1b-c*). DNA extraction was performed using the FastDNA^®^ SPIN Kit for Soil (MP Biomedicals, Solon, OH, USA) following the manufacturer’s protocol. The concentration and purity of the DNA extracts were measured using a TBS-380 fluorometer (Turner Biosystem, USA) and a NanoDrop 2000 spectrophotometer (Thermo Scientific, USA), respectively. The quality of the DNA extracts was checked using 1%-agarose gel electrophoresis. Paired-end sequencing was performed on the Illumina HiSeq XTen platform (Illumina, Inc., San Diego, CA, USA) at Majorbio Bio-Pharm Technology Co., Ltd., China. DNA extracts were fragmented to an average size of approximately 400 bp using a Covaris M220 ultrasonicator (Gene Company Limited, China). Paired-end libraries were constructed using the NEXTFLEX Rapid DNA-Seq Kit (Bioo Scientific, Austin, TX, USA), with adapters containing the full complement of sequencing primer hybridization sites ligated to the blunt end of fragments.

### Metagenome processing and MAG recovery

Raw reads were trimmed and quality-filtered using fastp v0.19.7^83^. The microbial diversity and its coverage based on quality-filtered raw metagenomic reads was estimated by Nonpareil v3.4.1^76^ in “kmer” mode. High-quality reads from individual samples were assembled into scaffolds using metaSPAdes version 3.15.5 (--k-list 21, 33, 55, 77, 99, and 127)^77^. Open-Reading Frames on scaffolds were predicted using prodigal (v2.6.3)^78^ in “meta” mode. MAGs were recovered from each assembly using the binning module (parameters: -metabat2 -maxbin2 -concoct) within metaWRAP v1.3 (metabat2 v2.12.1, MaxBin2 v.2.2.6, and concoct v0.4.0)^79–82^, which bins based on reads mapped only from the corresponding sample. Quality of genome bins (completeness and contamination) was evaluated using CheckM2 (v2.0.11)^83^. Binning results from multiple algorithms were integrated using DASTool v1.1.7 with default parameters^84^ to generate a non-redundant MAG set for each sample. The resulting bins were further refined using itBins (https://github.com/ProbstLab/itBins), and screened and cleaned for contaminant sequences with the NCBI Foreign Contamination Screen (FCS-GX v0.5.5)^85^. Finally, a total of 4,075 MAGs were obtained with completeness ≥50% and contamination <10%. Bins were further consolidated via dereplication at strain level using dRep (v.3.2.2) at 99% ANI^86^ resulting in 2,294 strain-cluster representative MAGs (strain-cluster rMAGs).

### Taxonomic classification of strain-cluster rMAGs and phylogenetic tree generation

Taxonomies were assigned to each strain-cluster rMAG using GTDB-Tk v2.1.1 (https://github.com/Ecogenomics/GTDB-Tk, database release 226)^87^. Phylogenetic relationships were inferred using GTDB-Tk (v2.1.1), which identifies conserved marker genes and constructs genome trees following the GTDB reference framework. The resulting GTDB-based phylogenetic tree was used to resolve the taxonomic placement of strain-cluster rMAGs. The phylogenetic tree was visualized using the Interactive Tree of Life (iTOL)^88^.

### Coverage estimation and metabolic potential prediction

Clean reads from each sample were mapped to strain-cluster rMAGs using strobealign (v0.13.0)^89^ in CoverM (v0.7.0; default parameters)^90^. The metabolic potential of strain-cluster rMAGs was determined using METABOLIC v4.0^91^. We constructed two databases, one containing 96 genes associated with DNA repair mechanisms and one containing 32 genes implicated in microbial responses to oxidative stress, with all genes being sourced from NCBI (https://www.ncbi.nlm.nih.gov/). Genes of strain-cluster rMAGs were annotated against these databases using DIAMOND BLAST^92^ with an E-value threshold of 1e^-5^ and the parameter -k 1 to retain only the top scoring hit. To ensure the specificity of these annotations, a reverse BLAST was subsequently conducted by aligning database hits back against the original amino acid sequences of the corresponding strain-cluster rMAGs using the same E-value threshold. Only reciprocal hits that aligned to the same ORFs as the original query were retained, ensuring one-to-one correspondence. Cases with multiple high-scoring matches were excluded to avoid ambiguous annotations. Final hits were filtered based on a similarity threshold of >50%, ensuring high-confidence functional assignments. These curated datasets facilitated a refined analysis of microbial resilience and adaptation strategies in diverse environmental contexts.

### Community profiling based on the *rpS3* marker gene

ORFs representing the *rpS3* marker gene were identified using hmmsearch^93^ against custom Hidden Markov Models (HMMs) for Archaea and Bacteria (E-value 1e^-28^; https://github.com/AJProbst/rpS3_trckr). Sequences were pooled and clustered at 99% similarity using vsearch^94^, representing species-level clusters^95^. Representative sequences were selected following a tiered approach: (1) centroids that could be extended by 1000 bp in both directions, (2) non-centroids that were similarly extendable, and (3) the longest available sequence when neither of the first two criteria were met. To improve abundance estimates, *rpS3* gene sequences of representative *rpS3* genes were extended by up to 1,000 bp in both directions on their scaffold of origin if possible. This was done as mapping on just the approximately 1,000 bp long *rpS3* gene results in poor mapping on the gene edges and this has progressively more impact when the total length of the sequence is short. The resulting *rpS3* gene sequences, ideally extended in both directions by 1,000 bp, are henceforth called *rpS3*-extended. The workflow for clustering, extracting, and calculating coverage is available at https://github.com/ProbstLab/publication-associated_scripts/tree/main/FigueroaGonzalez_Bornemann_etal. The *rpS3* genes in *rpS3*-extended sequences (in amino acid format) were taxonomically classified using DIAMOND^92^ (E-value ≤ 1e^−5^) against the GTDB r226 protein database, assigning the best hit to be the taxonomy. If no hit meeting the search criteria was found, “Unclassified” was assigned. Abundance was calculated by mapping reads of all metagenomic samples to *rpS3*-extended sequences and calculating the average coverage per sample. Additionally, the breadth, i.e., the number of nucleotide positions in *rpS3*-extended having a coverage of at least 1, was determined. The coverage of an *rpS3*-extended sequence in a given sample was set to zero if the breadth of the sequence in that sample was less than 95%, indicating spurious mapping instead of actual presence of the *rpS3*-extended-represented species. Relative coverage values were transformed into percentages (%).

### RNA extraction, sequencing, and metatranscriptomics

Total RNA was isolated and purified using Soil RNA Extraction Kit (Majorbio, Shanghai, China) following the manufacturer’s procedure. The integrity and quantity of the extracted RNA were measured with a NanoDrop 2000 spectrophotometer (Thermo Scientific, MA, USA) and an Agilent 5300 Bioanalyzer (Agilent Technologies, Palo Alto, CA, USA). The RNA was subjected to standard Illumina library preparation using the Illumina® Stranded mRNA Prep, Ligation kit (Illumina, San Diego, CA, USA), and rRNA was depleted using a RiboCop rRNA Depletion Kit for Mixed Bacterial Samples (Lexogen, USA). Depleted RNA was extracted from biomass retained on 0.22 μm sterile cellulose nitrate membranes from 60 groundwater samples (same to metagenomic groundwater samples) and finally library preparation of RNA extraction from 45 groundwater samples were successfully constructed (Figure *1c*). Sequencing was performed on an Illumina Novaseq6000 sequencer (Illumina, San Diego, CA, USA) with paired-end sequencing method at Majorbio Bio-Pharm Technology Co., Ltd. (Shanghai, China). Quality control of the raw data generated by sequencing was carried out with fastp v0.19.7^96^. SortMeRNA (v4.3.6) was used to remove non-coding RNA sequences (tRNA, tmRNA, 5S, 16S, 18S, 23S, and 28S)^97^. The proportion of reads removed per sample is summarized in Supplementary Table *S3* and Figure *S20*, providing an overview of rRNA/tRNA content prior to filtering.

### Gene expression dynamics in metatranscriptomes

InStrain (v1.8.1)^98^ with the parameter (--min_cov 0) was used on strain-cluster rMAGs recovered from the same sample to get the coverage of each gene using the clean reads from each metagenome and each metatranscriptome. The gene coverage in each strain-cluster rMAG was received by using the cluster database generated when dereplicate genomes at 99% ANI using dRep (v3.2.2)^86^. The mean coverage of 30S ribosomal proteins were used to calculate the normalization factor of each gene from the strain-cluster rMAGs in each metatranscriptome sample. The genes with normalization expression level greater than two were defined as overexpressed genes. The average expression level of each gene in each sample was calculated and subjected to Principal Coordinates Analysis (PCoA) based on Bray-Curtis dissimilarity. Average gene expression levels were also used to calculate Spearman correlations with TU and radioactivity. In addition, a presence-absence matrix was constructed using gene annotation profiles predicted from metabolic potential analyses.

### Species-level clustering and calculation of evolutionary metrics

Filtered reads from each metagenome and metatranscriptome were mapped to the cleaned MAGs output from FCS-GX (v0.5.5) of the same sample using strobealign (v0.13.0)^89^ in CoverM (v0.7.0; default parameters)^90^. Open reading frames (ORFs) of each MAG were predicted using Prodigal (v2.6.3; –p single)^78^ and used as input for inStrain (v1.8.1)^98^. Population statistics and nucleotide metrics, including nucleotide diversity (SNVs/kbp) and nonsynonymous to synonymous mutation ratios (pN/pS) were calculated from these mappings using the profile module of the inStrain (v1.8.1; default parameters)^98^ at genome and gene levels. The 4075 MAGs reconstructed from all individual assemblies (n = 60) were combined and dereplicated for species-level clustering using dRep (v3.2.2)^86^ at 95% ANI. A total of 1752 MAGs with the highest genome quality from each species cluster were designated as the representative species (species-cluster rMAGs). The species-cluster rMAGs with over three members in the cluster database generated when dereplicate genomes at 95% ANI were selected for the final calculation of microdiversity metrics such as coverage, breadth, SNPs density, nucleotide diversity, and pN/pS ratios.

For SNP calling, the number of quality-filtered reads mapping to the position had to be at least 5× coverage and 5% SNP frequency, and the variant based on reads had to be lower than the expected sequencing error rate (1e^−6^). Genome-level evolutionary metrics based on both metagenomic and metatranscriptomic data from mining sites were compared with those from background aquifer samples at the same location (Table *S4-S5*). To assess the genomic impact of radiochemical stress, four species-cluster rMAGs showing increased nucleotide diversity and detected in at least two groups (BG, MT, and PM) with a minimum of three samples per group were selected. Gene annotations for the four species-cluster rMAGs were generated using MetaCerberus v1.3.1^99^. All member genomes within the four species-cluster were used to identify representative genes using mmseqs2 v17.b804f^100^, from which median nucleotide diversity was calculated from both metagenomic and metatranscriptomic datasets across the BG, MT, and PM groups (*Table S6-S7*).

### Statistical analyses and data visualization

Spearman correlation analyses were performed in the R platform (http://cran.r-project.org), with p-values adjusted using the Benjamini-Hochberg procedure at a minimum significance threshold of 0.05^101^. Co-occurrence networks were constructed using the *igraph* package with default parameters^102^. Only Spearman correlations with p-values < 0.001 and |ρ|> 0.6 between pairs of rMAGs were considered statistically significant, contributing to the construction of a reliable network. The resulting correlation network was imported into Gephi (v0.9.2) for visualization^103^. The Spearman correlations between over-expressed genes from strain-cluster rMAGs and hydrogeochemical variables were visualized using the *pheatmap* package in R (v4.4.2)^104^. To assess sample similarity and identify factors shaping community clustering, Principal Coordinates Analysis (PCoA) and Non-metric Multidimensional Scaling (NMDS) were performed based on Bray–Curtis dissimilarity of microbial community composition. Relationships between geochemical parameters and biological features—including community dissimilarities, gene expression levels, nucleotide diversity, and pN/pS ratios—were further evaluated using *Bioenv* analysis (*vegan* package, R v4.4.2)^105^, which identifies the combination of environmental variables best explaining the observed biological variation.

## ACKNOWLEDGEMENTS

This study was supported by the National Natural Science Foundation of China grant nos. 42130509 (to H.G.) and 42072273 (to W.X.), the Deep Earth Probe and Mineral Resources Exploration-National Science and Technology Major Project grant no. 2024ZD1000601 (to W.X.), the Fundamental Research Funds for the Central Universities grant no. 590223006 (to W.X.) and the 111 Project grant no. B20010 (to H.G.); and by the European Research Council Synergy Grant “Archean Park” under grant no. 101118631 (to A.J.P.) and the German Research Foundation grant no. PR 1603/4-1 (to A.J.P.). We thank Prof. Rizlan Bernier-Latmani (Environmental Microbiology Laboratory, École Polytechnique Fédérale de Lausanne, Switzerland) for valuable comments on the manuscript and for providing access to laboratory resources.

## Ethics Statement

All sampling and sample collection were performed with permission from the local environmental authorities and in compliance with all applicable regulations governing environmental research in China. No human subjects or animal samples were involved in this study.

## CODE AVAILABILITY STATEMENT

All code used in this manuscript is publicly available and referenced in the Materials and Methods section.

