## Supplementary material for "Uranium mining fuels evolution in deep groundwater microbiomes": Supplymentary_material

**Figure S1 | *Geochemical profiles of groundwater across neutral-pH ISL phases.*** *Boxplots show geochemical parameters through the stage of neutral-pH ISL, including pre-mining (BG), mining (MT), and post-mining (PM). Sampling groups include: Site 1 (unmined aquifer, S1_BG); Site 2 (unmined aquifer, S2_BG; recent ISL at 0.25-1 years: S2_MT0.25 and S2_MT01); Site 3a (unmined aquifer, S3a_BG; intermediate ISL, 4-6 years: S3a_MT04, S3a_MT05, and S3a_MT06); and Site 3b (unmined aquifer, S3b_BG; long-term ISL, 15-20 years: S3b_MT15 and S3b_MT20; and decommissioned ISL under iron-reducing, S3b_PMI, mixed iron- and sulfate-reducing, S3b_PMIS, and sulfate-reducing, S3b_PMS conditions). Panels (a – h) represent sulfide, sulfate, sulfate/Cl, Fe(II), ammonium, nitrite, nitrate, and total Fe, respectively. Colored backgrounds denote different ISL phases. Colors indicate study sites: light blue (Site 1), light green (Site 2), medium green (Site 3a), and beige (Site 3b). Statistical significance was determined by ANOVA with Tukey's or Kruskal–Wallis with Dunn's test (BH-corrected), depending on data normality.*

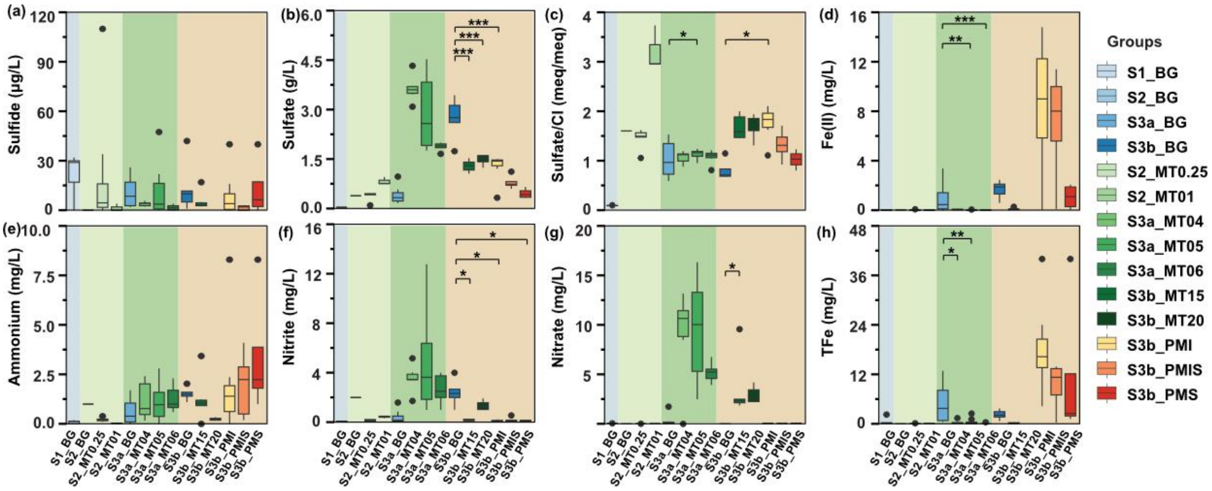

**Figure S2 | Spearman correlation matrix among geochemical parameters in groundwater samples.** Pairwise Spearman correlation coefficients are visualized as a color-coded heatmap. Statistical significance is denoted by asterisks (* $p < 0.05$, ** $p < 0.01$, *** $p < 0.001$). Variables include radioactive indicators (gross alpha and beta activity), physicochemical parameters (e.g., pH, Eh, TDS, and DOC), and redox-sensitive species (Fe(II), sulfide, ammonium, and nitrate). The heatmap highlights strong covariation among multiple geochemical stressors in uranium ISL-impacted groundwater systems.

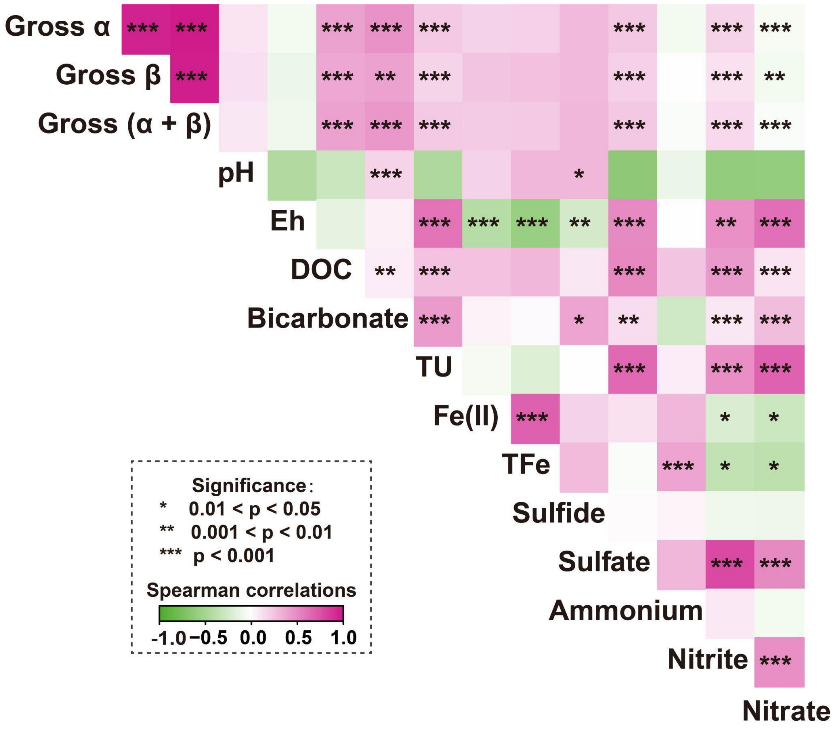

**Figure S3 | Nonpareil analysis of average sequencing coverage and estimated sequencing effort.**

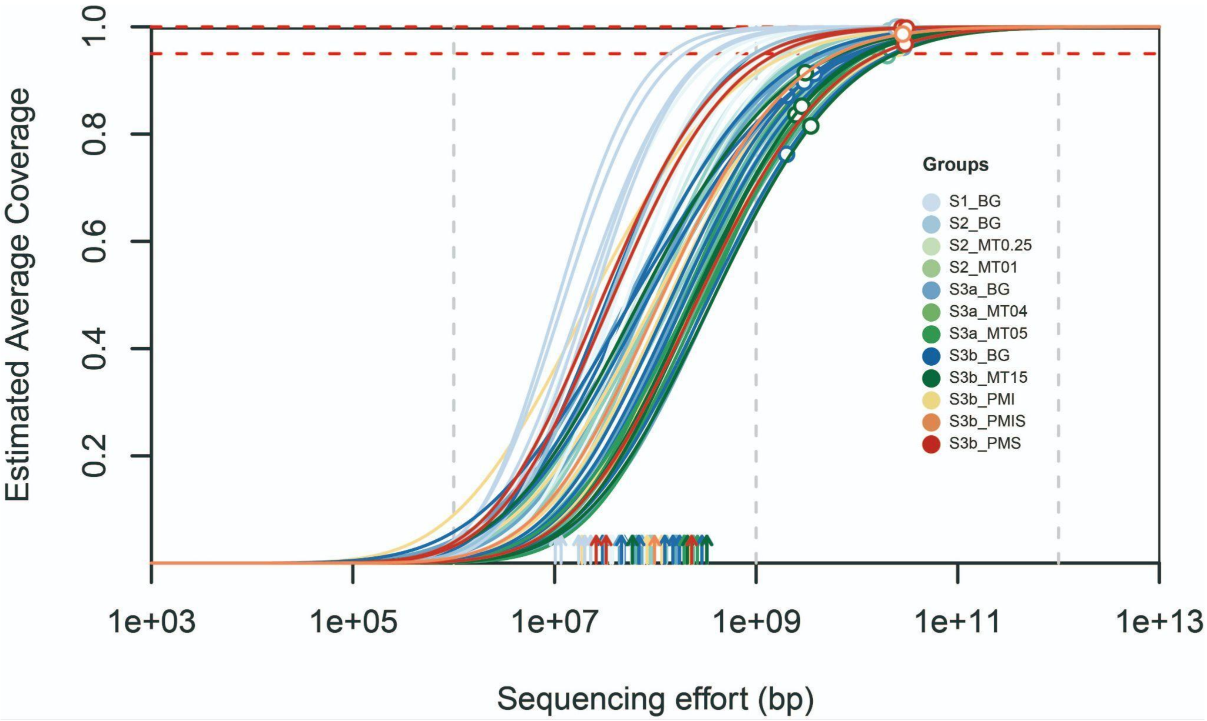

**Figure S4 | Comparison of taxonomic resolution between metagenomic (metaG) and rpS3 gene-based profiling across taxonomic ranks**. Bar chart showing the number of taxa ($log_{10}$ scale) identified at each taxonomic level (phylum to genus) using genome-resolved metagenomic data (blue) versus rpS3 gene-based analysis (orange).

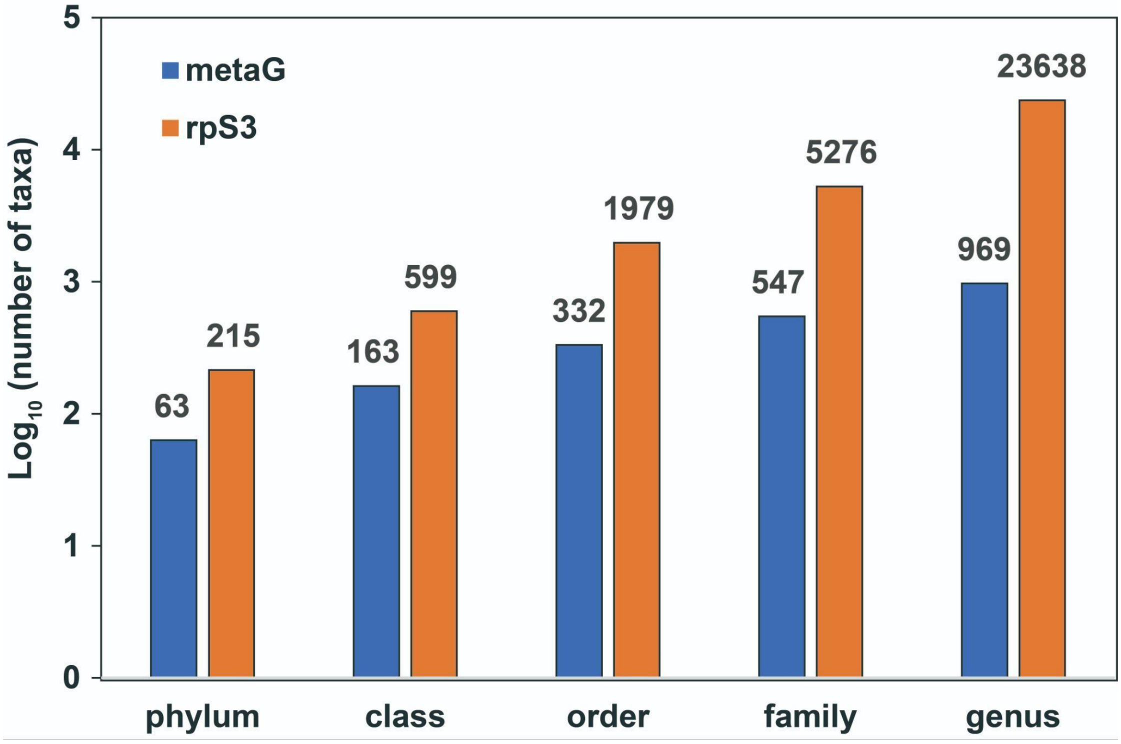

**Figure S5 | Alpha diversity patterns of groundwater microbial communities based on *rpS3* gene profiles and their correlations with environmental factors.** (a–c) Boxplots of alpha diversity indices across sites and treatment stages: (a) Shannon index, (b) Simpson index, and (c) Inverse Simpson index. Differences between groups were assessed using ANOVA followed by Tukey's post-hoc test and indicated by asterisks (*p < 0.05, **p < 0.01, ***p < 0.001); (d) Spearman correlation heatmap showing associations between environmental variables and alpha diversity metrics. Blue indicates negative correlations, red indicates positive correlations. Dendrogram shows hierarchical clustering of environmental parameters based on correlation patterns.

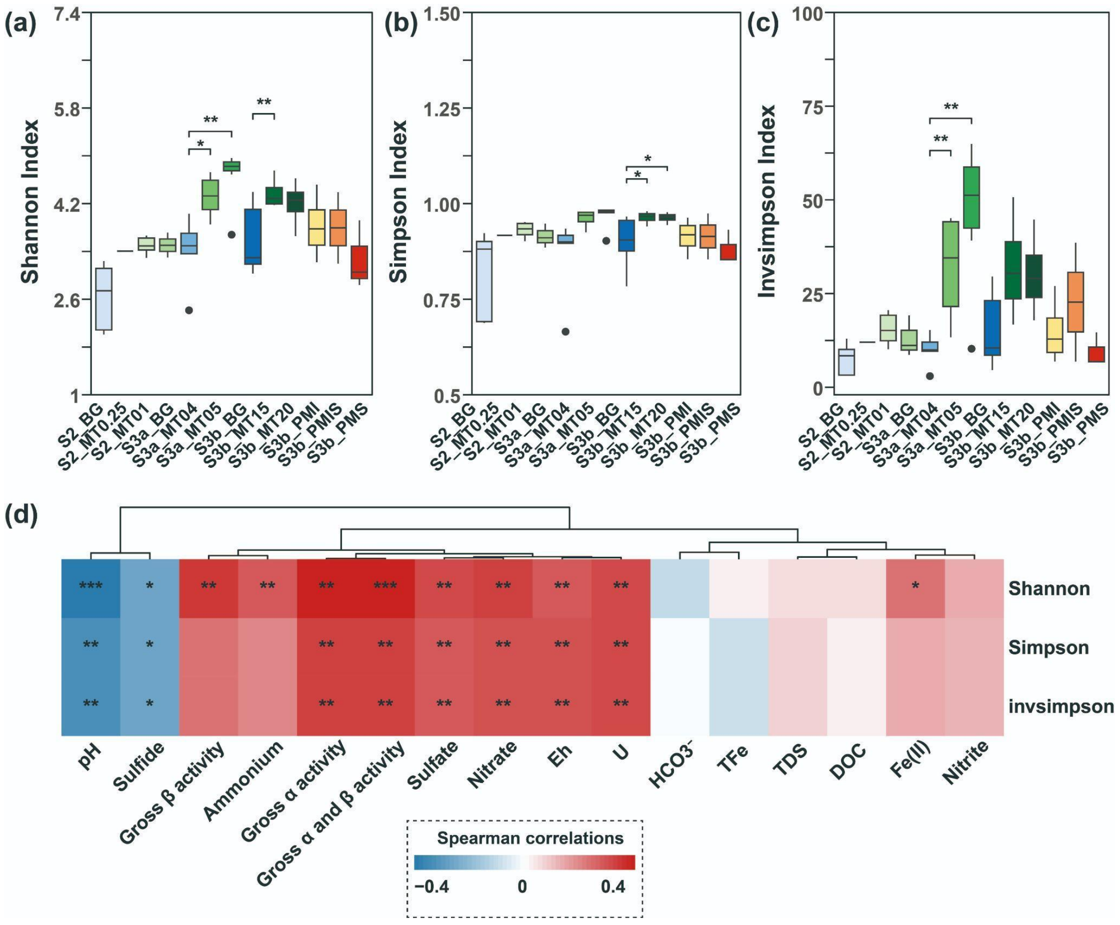

**Figure S6 | Beta diversity of groundwater microbial communities based on *rpS3* gene profiles.** (a) Principal Coordinates Analysis (PCoA) of microbial community composition (based on percent relative abundance of *rpS3* gene sequences across samples), constrained by groundwater geochemical variables. Vectors represent environmental parameters significantly correlated with community variation, including pH, sulfide, total Fe (TFe), Fe(II), redox potential (Eh), bicarbonate ($HCO_3^-$), and gross alpha and beta radioactivity; (b) Non-metric multidimensional scaling (NMDS) ordination of *rpS3*-based community dissimilarity, overlaid with uranium concentration (U, mg/L) contours. Point colors correspond to sampling sites and ISL stages, as indicated in the legend. Stress value indicates ordination quality (stress = 0.176).

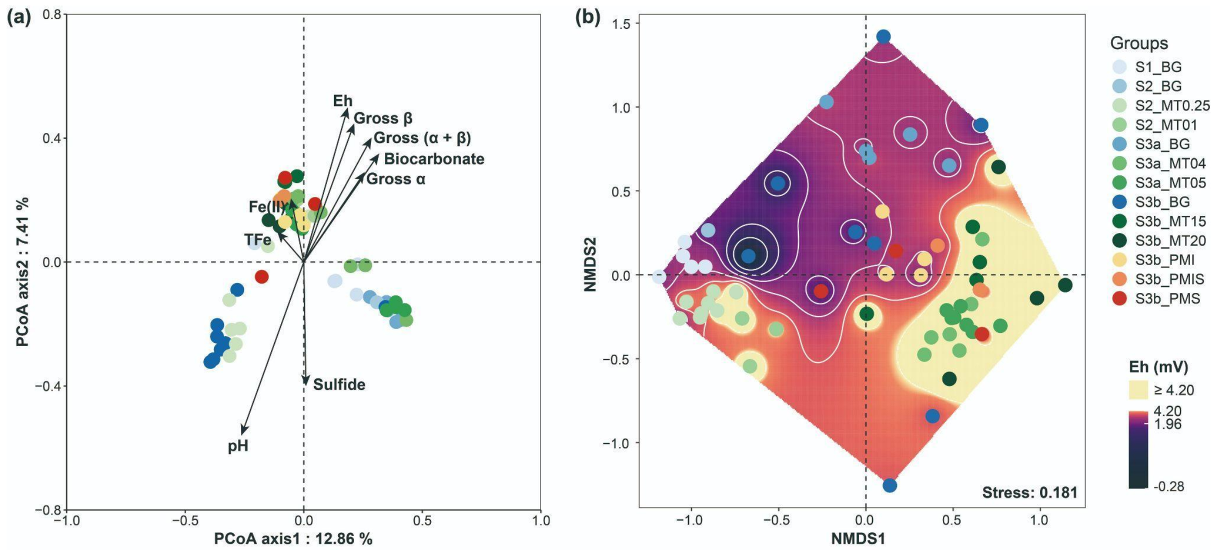

**Figure S7 | Percentages of metagenomic reconstuctued to rpS3-encoded strain-cluster rMAGs across groundwater sites and ISL stages**. Group colors correspond to sampling sites and ISL stages as indicated in the legend.

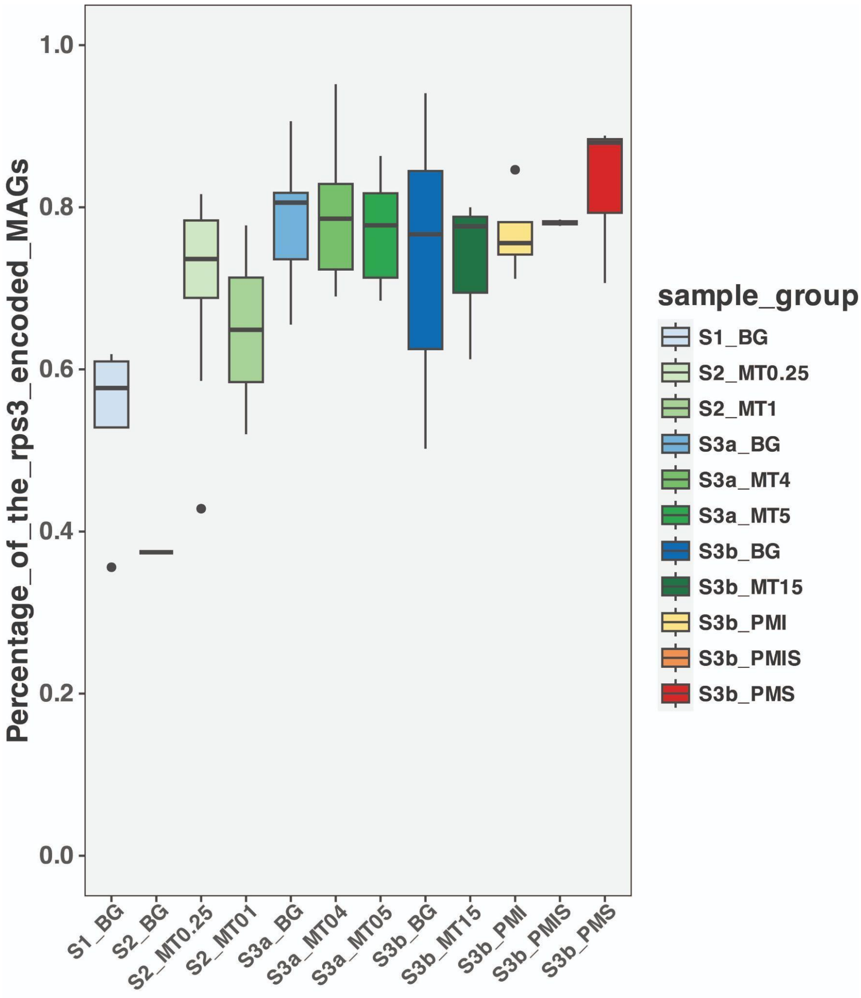

**Figure S8 | Distribution of strain-cluster rMAGs across groundwater samples in different ISL stages.** Heatmap shows the Z-score normalized relative abundance (RPKM) of strain-cluster rMAGs across groundwater samples in different ISL stages.

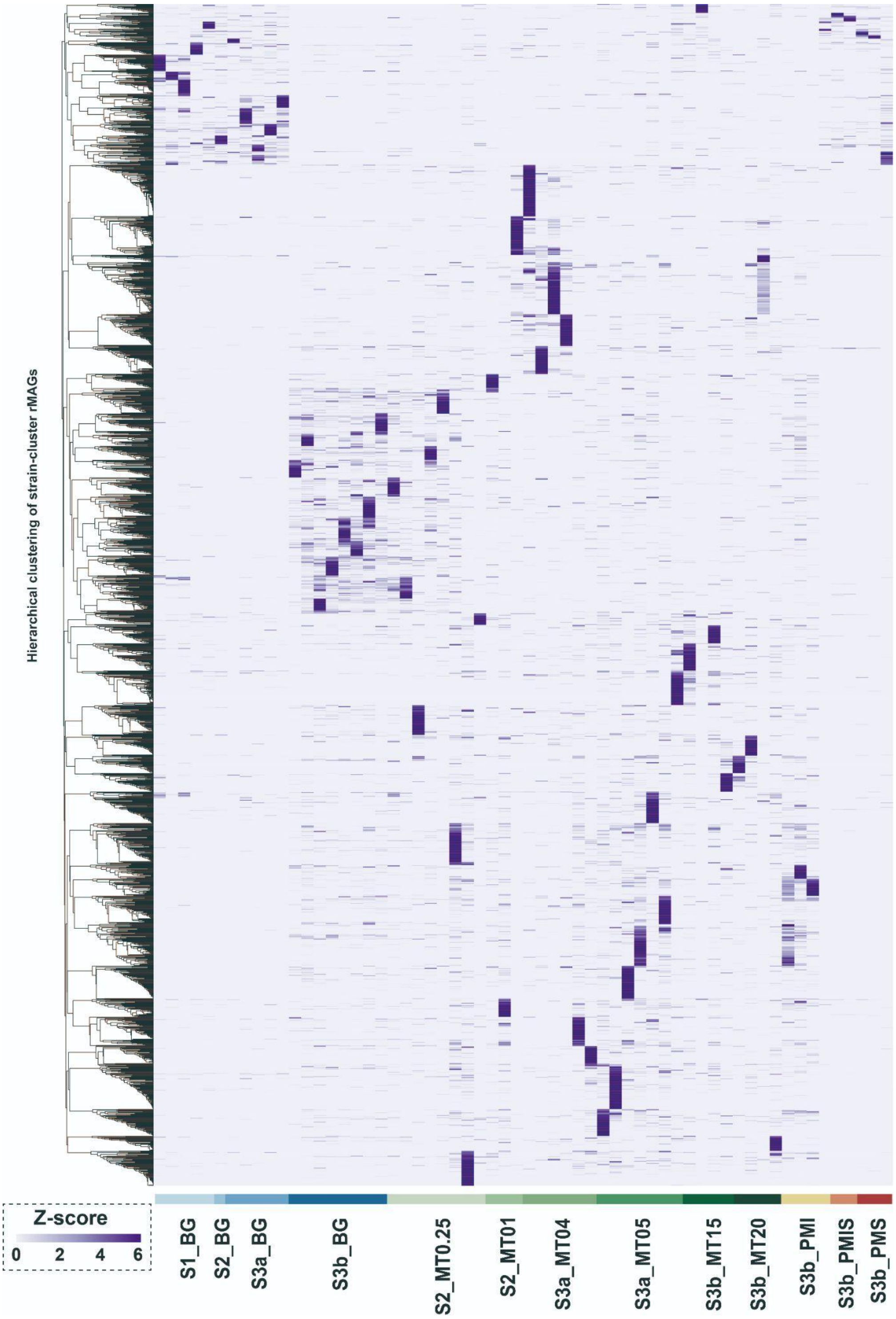

**Figure S9 | Correlations of overexpressed genes with hydrogeochemical parameters.** Heatmap showing significant positive Spearman correlations ($p < 0.05$, $\rho > 0$) between overexpressed genes and hydrogeochemical parameters. Each gene and parameter is color-coded by functional category or chemical group. The accompanying bar chart indicates the number of strain-cluster rMAGs encoding the corresponding overexpressed genes.

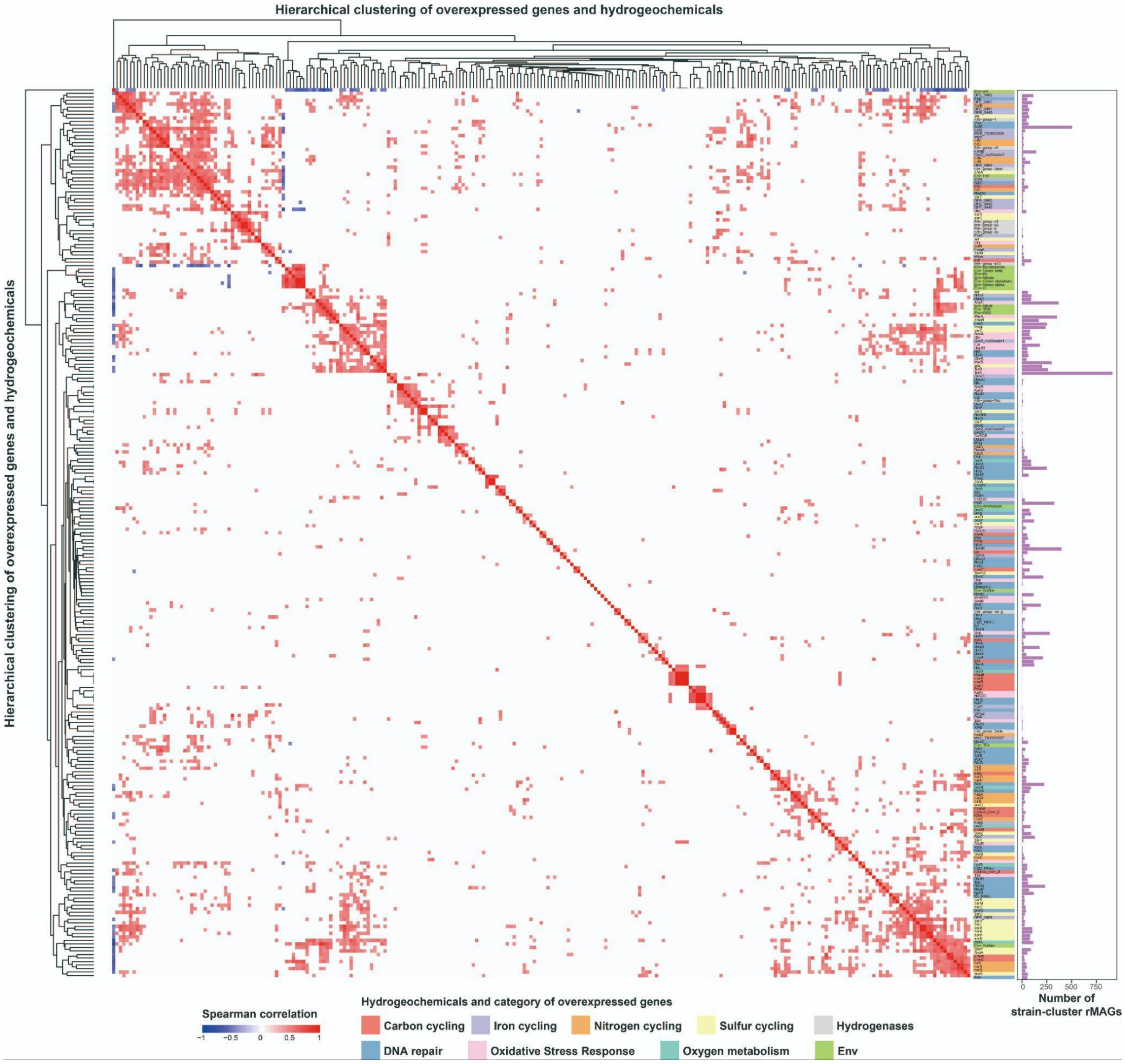

**Figure S10 | Transcriptional enrichment of oxidative stress–response genes in strain-cluster rMAGs.** Boxplots show $log_2$-transformed expression levels of gene groups associated with (A) lipid repair, (B) protein repair, (C) stress-response regulators, and (D) reactive oxygen species (ROS) scavenging across different groundwater treatment stages. Overexpression was determined based on metatranscriptomic data relative to background expression baselines. Groups are color-coded by sampling sites and ISL stages (see legend).

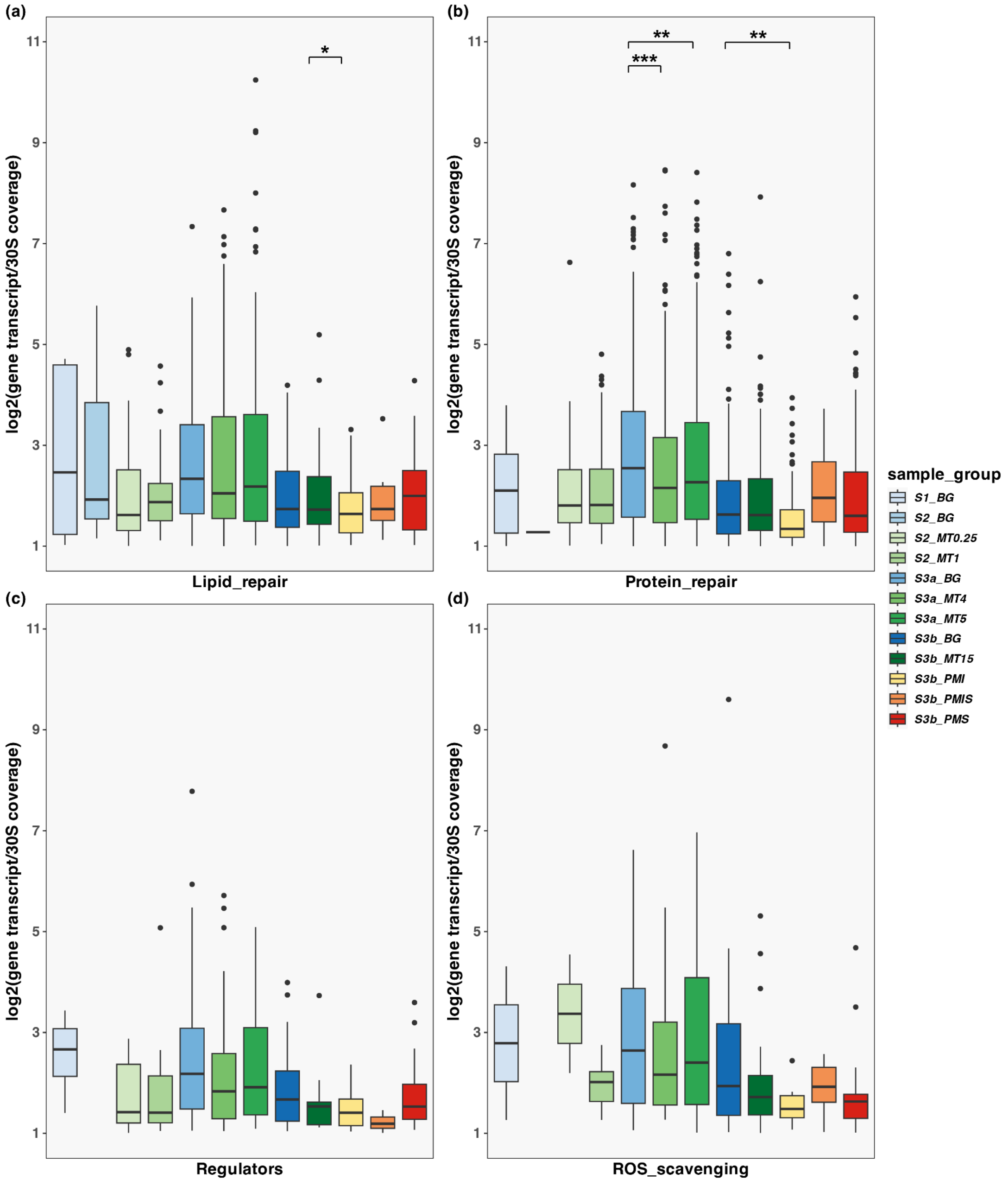

**Figure S11 | Metatranscriptomic overexpression profiles of metabolic pathways in strain-cluster rMAGs across groundwater sites.** Boxplots show $log_2$-transformed expression levels of genes grouped by metabolic functions across background (BG), mining (MT), and post-mining (PM) stages. Functional categories include (top row) $C_1$ metabolism, carbon fixation, and fermentation; (middle row) methane metabolism, hydrogenases, and oxygen metabolism; and (bottom row) nitrogen, iron, and sulfur cycling. Overexpressed genes were identified from metatranscriptomic data relative to baseline background expression. Site groups are color-coded as shown in the legend.

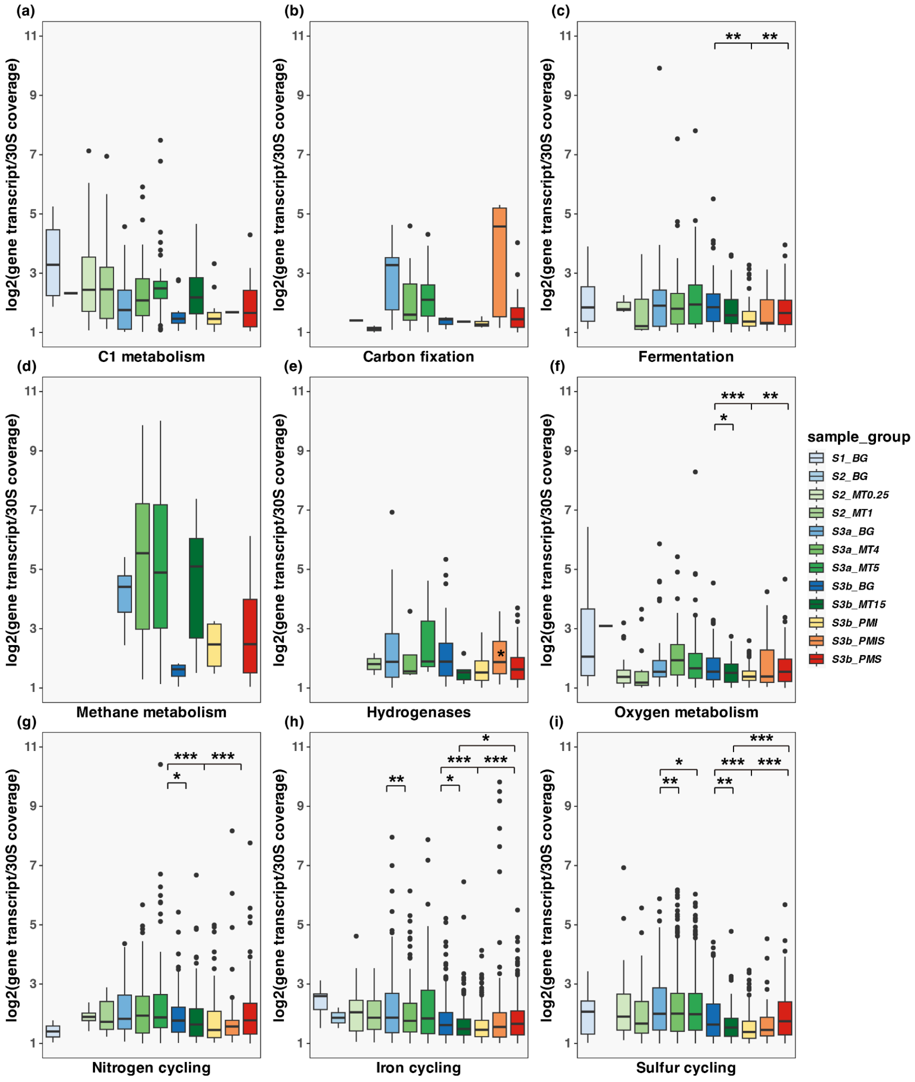

**Figure S12 | Transcriptional activity of nitrogen cycling pathways in strain-cluster rMAGs across groundwater samples.** Boxplots display $log_2$-transformed expression levels of genes involved in nitrogen transformations across background (BG), mining (MT), and post-mining (PM) stages. Functional groups include nitrite reduction to ammonia, $N_2$ fixation, nitric oxide reduction, nitrite reduction, nitrate reduction, ammonia oxidation, nitrite oxidation, anammox, and nitrous oxide reduction. Expression levels were derived from metatranscriptomic datasets and grouped by site category, with colors corresponding to sampling sites and ISL stages.

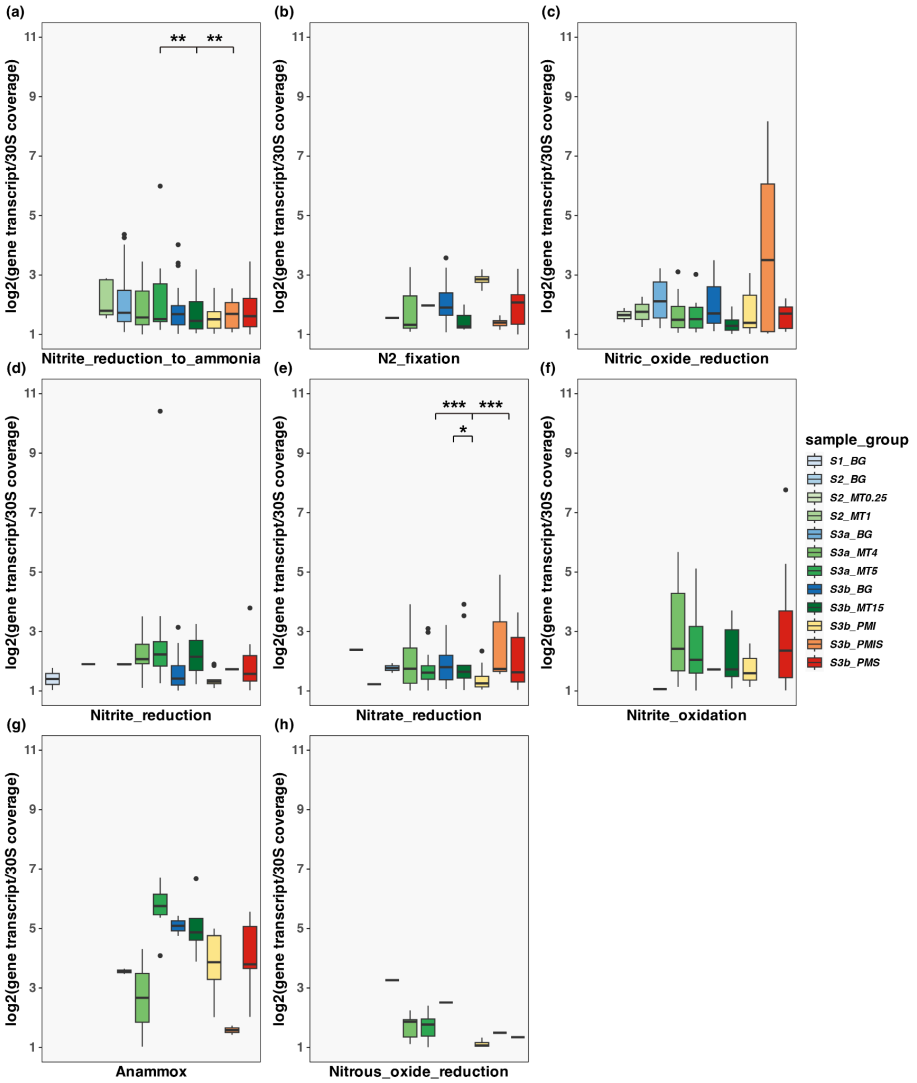

**Figure S13 Transcriptional expression of iron redox cycling genes in strain-cluster rMAGs across groundwater treatment stages |** Boxplots show $log_2$-transformed expression levels of gene groups associated with (A) iron reduction and (B) iron oxidation across background (BG), mining (MT), and post-mining (PM) sites. Expression values were derived from metatranscriptomic data and grouped by sampling sites and ISL stages as indicated in the legend.

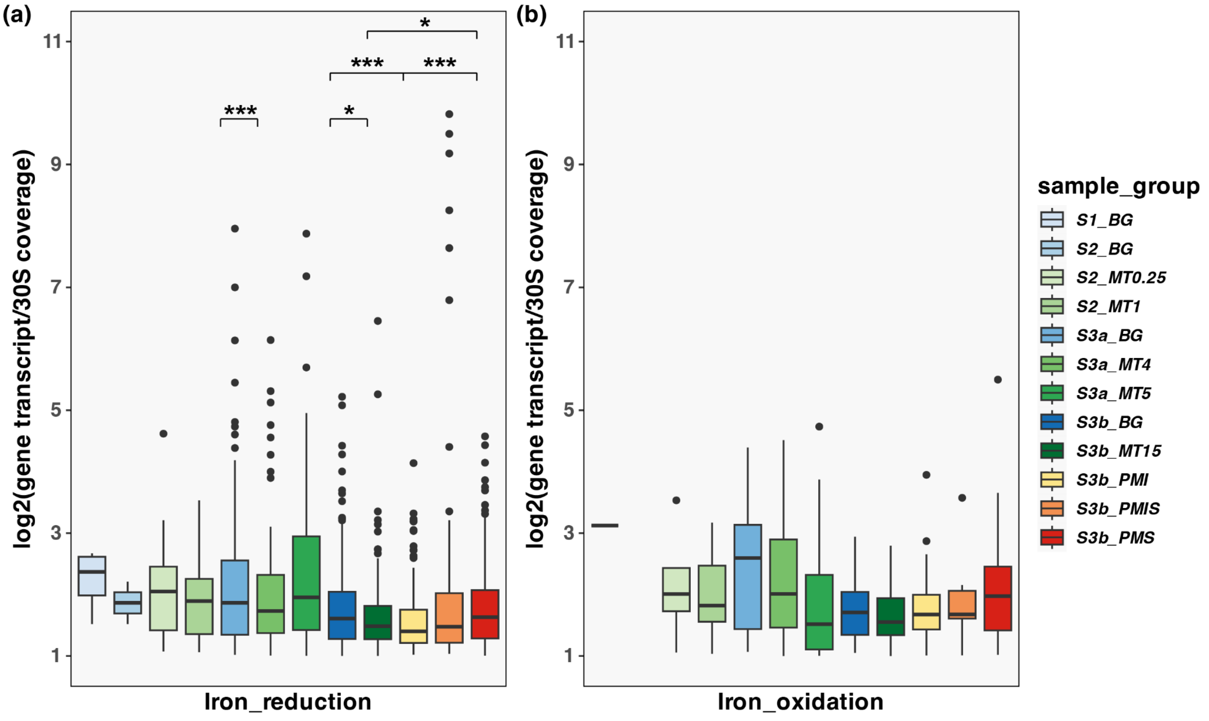

**Figure S14 | Transcriptional expression of sulfur metabolism pathways in strain-cluster rMAGs across groundwater samples.** Boxplots show $log_2$-transformed expression levels of genes associated with sulfur transformations, including sulfur oxidation, sulfide oxidation, thiosulfate oxidation, sulfate reduction, sulfite reduction, thiosulfate disproportionation, dissimilatory sulfur metabolism, DMSO metabolism, and sulfur-related amino acid metabolism. Expression values are based on metatranscriptomic profiles and grouped by sampling sites and ISL stages as indicated in the color-coded legend.

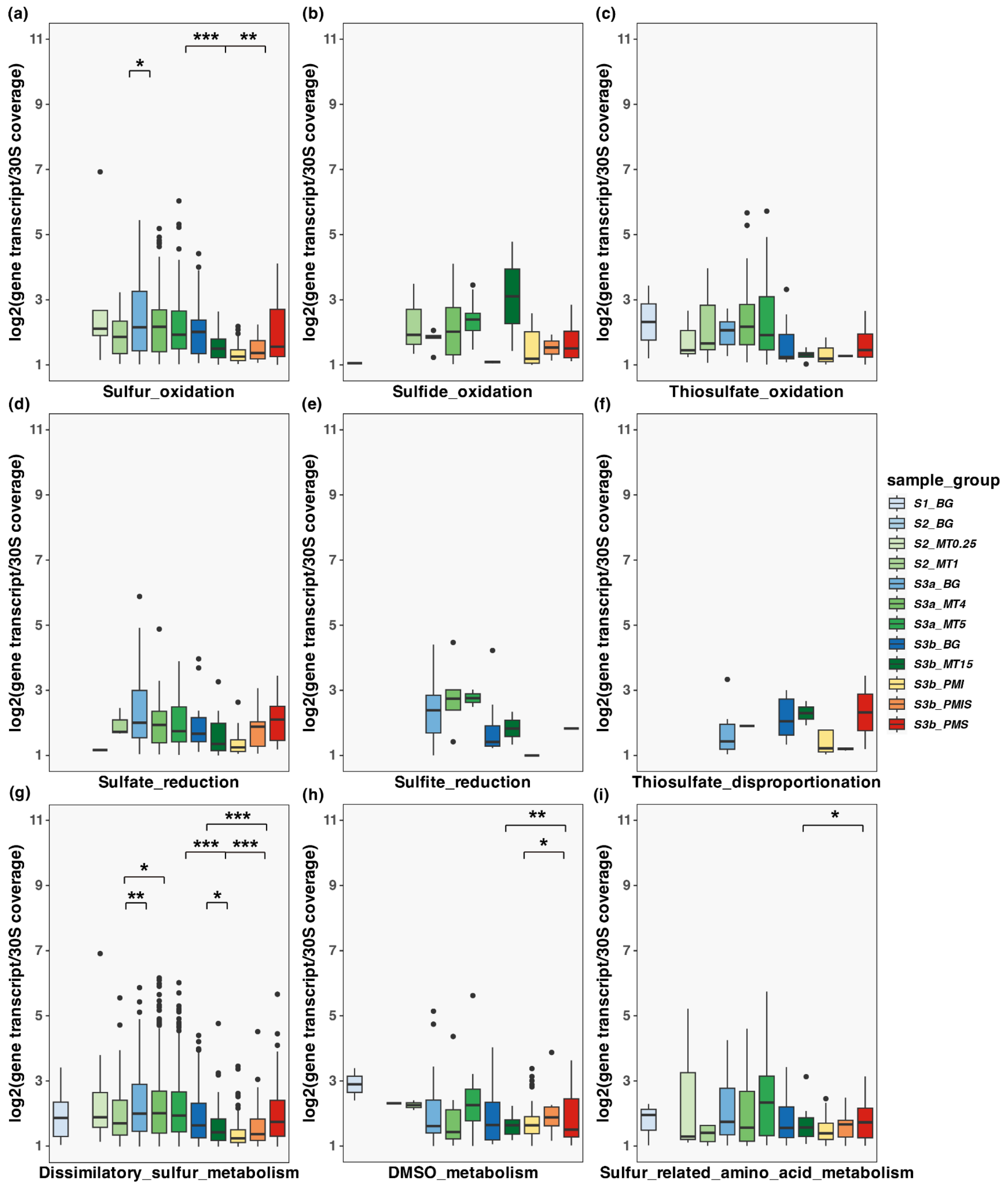

**Figure S15 | Transcriptional profiles of DNA repair pathways in strain-cluster rMAGs across groundwater samples under varying uranium ISL conditions.** Boxplots display $log_2$-transformed expression levels of genes involved in distinct DNA repair mechanisms, including base excision repair, mismatch repair, homologous recombination repair, nucleotide excision repair, direct reversal repair, DNA damage signaling, translesion synthesis, error-prone double-strand break (DSB) repair, and non-homologous end joining. Expression data were derived from metatranscriptomic datasets and grouped by sampling sites and ISL stages as indicated in the legend.

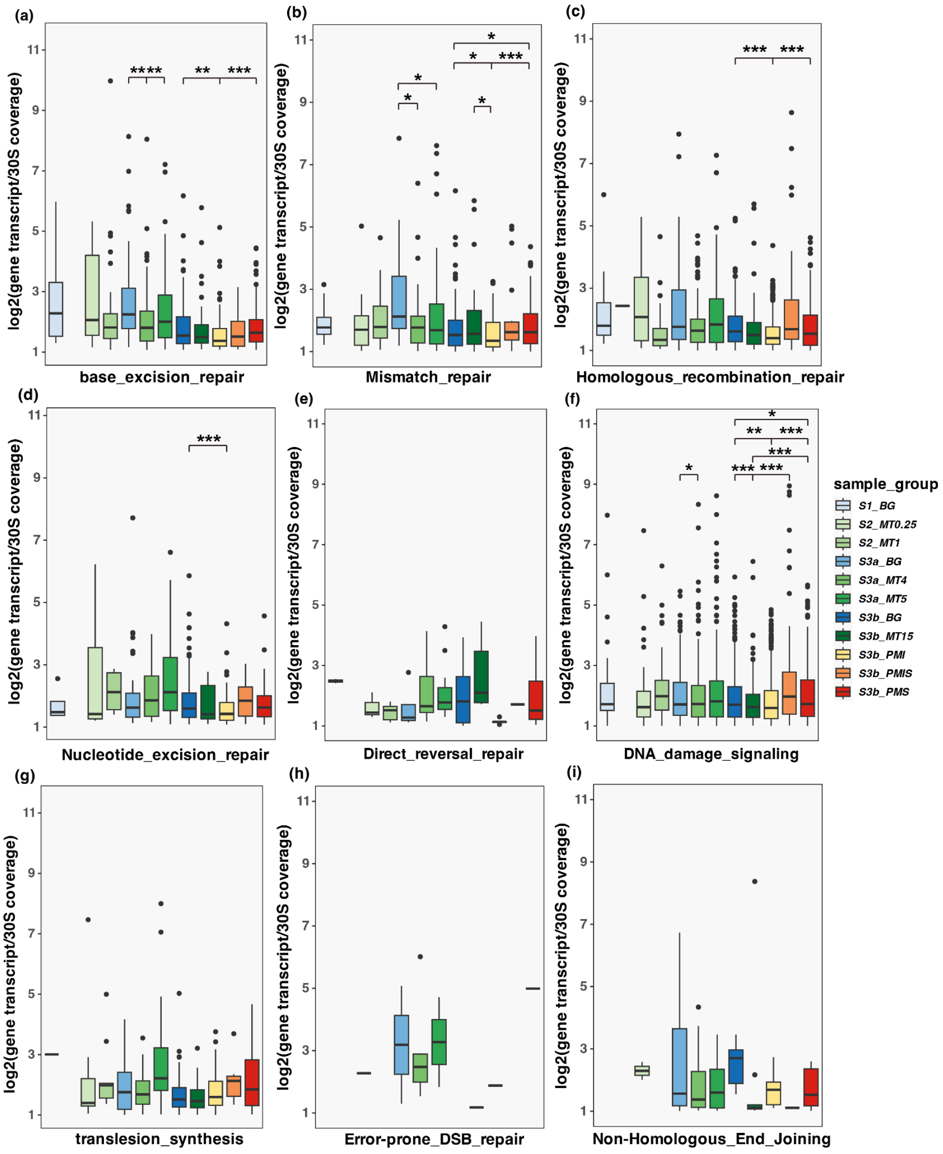

**Figure S16 | Expression patterns of key DNA repair genes in strain-cluster rMAGs across mining stages.** Boxplots show the normalized expression ($log_2$-transformed gene coverage normalized to 30S ribosomal protein coverage) of representative genes involved in multiple DNA repair pathways across aquifers impacted by uranium in situ leaching (ISL). Pathways include the SOS response (*LexA*, *RecA*, *AidB*), homologous recombination (*Ssb*, *RecQ*, *RuvC*, *RecG*), base excision repair (*ExoA*, *Fpg*), nucleotide excision repair (*UvrD*, *PolII*), mismatch repair (*MutL*), translesion synthesis (DinB), and double-strand break repair (*LigD_NHEJ*, *DNA_ligase_ligAB*). Groups are color-coded by mining stage: background (blue), mining (green), and post-mining (orange/red).

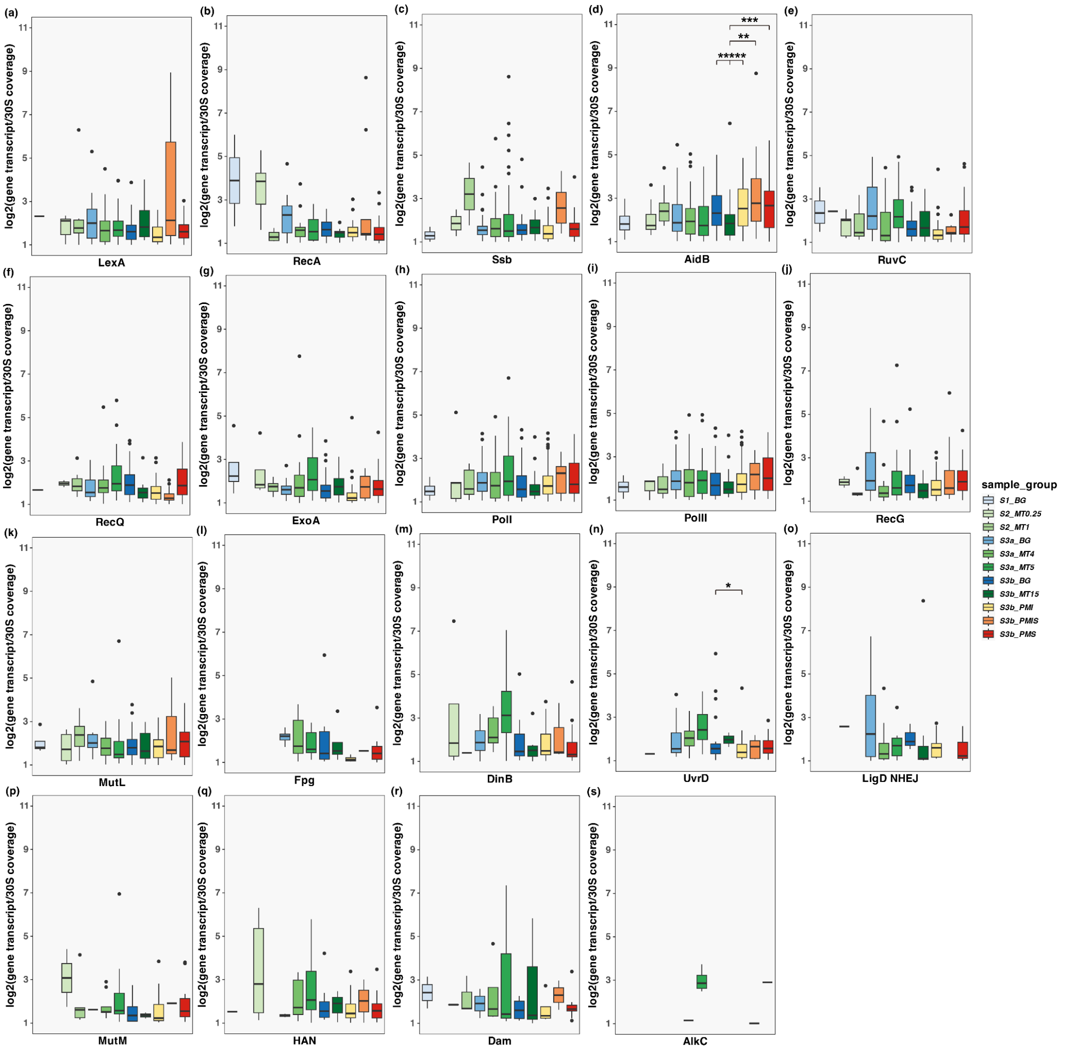

**Figure S17 | Expression patterns of C-H-O-N-S-Fe metabolic genes showing positive Spearman correlations with total dissolved uranium (TU), gross α, β, and (α + β) radioactivity.**

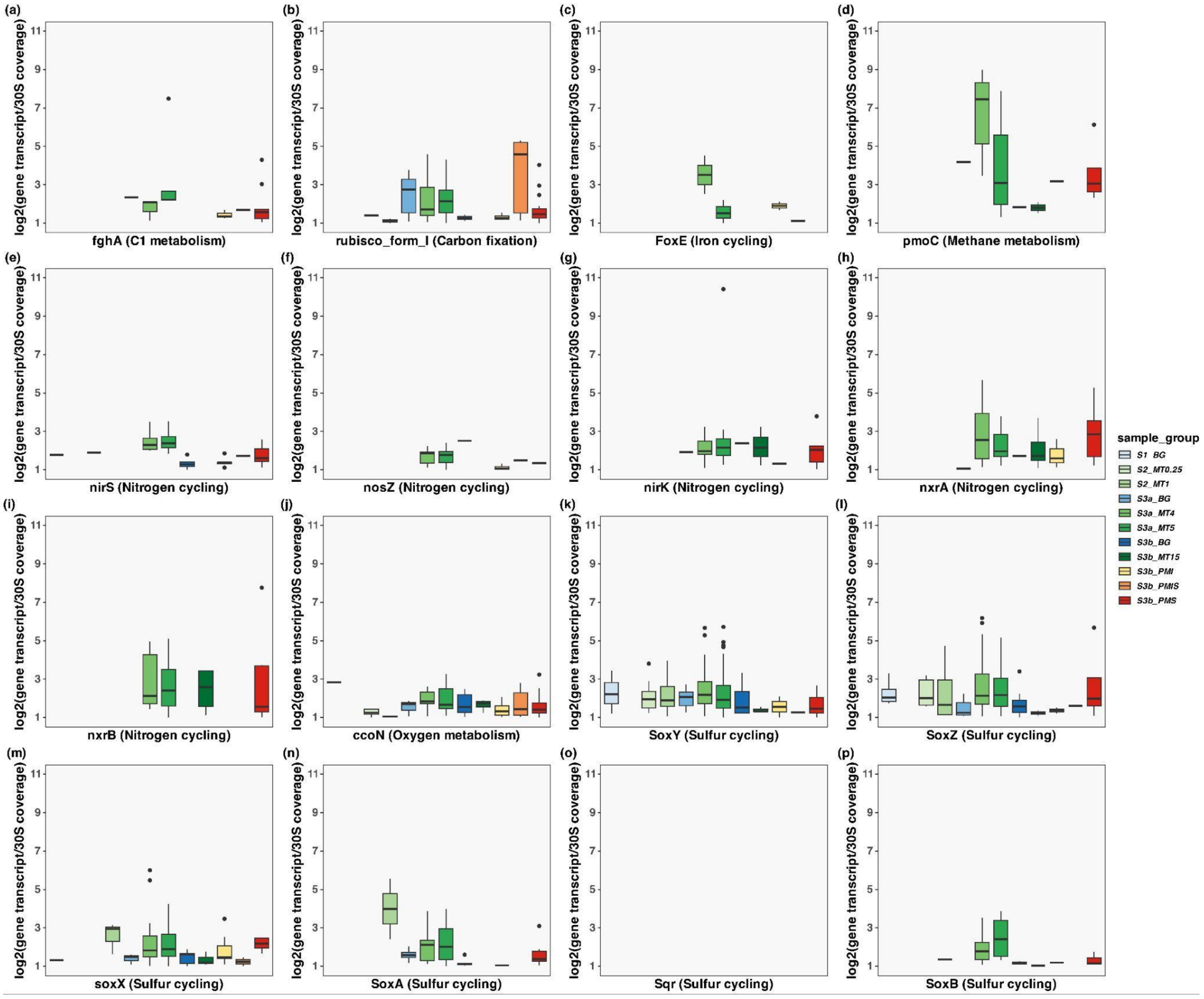

**Figure S18 | Transcriptional enrichment and functional partitioning of genes in species-cluster rMAGs**. (a) Flow diagrams illustrate the all genes with measurable nucleotide diversity, nucleotide diversity of genes in metagenomes and metatranscriptomes, and nucleotide diversity of overexpressed genes involved in biogeochemical cycling (C-H-O-N-S-Fe cycles); (b) Flow diagrams illustrate the all genes with non-synonymous SNPs, number of genes showing different changing patterns of pN/pS ratios in metagenomic and metatranscriptomic datasets, and number of overexpressed genes involved in biogeochemical cycling (C-H-O-N-S-Fe cycles) showing different changing patterns of pN/pS ratios in metagenomic and metatranscriptomic datasets.

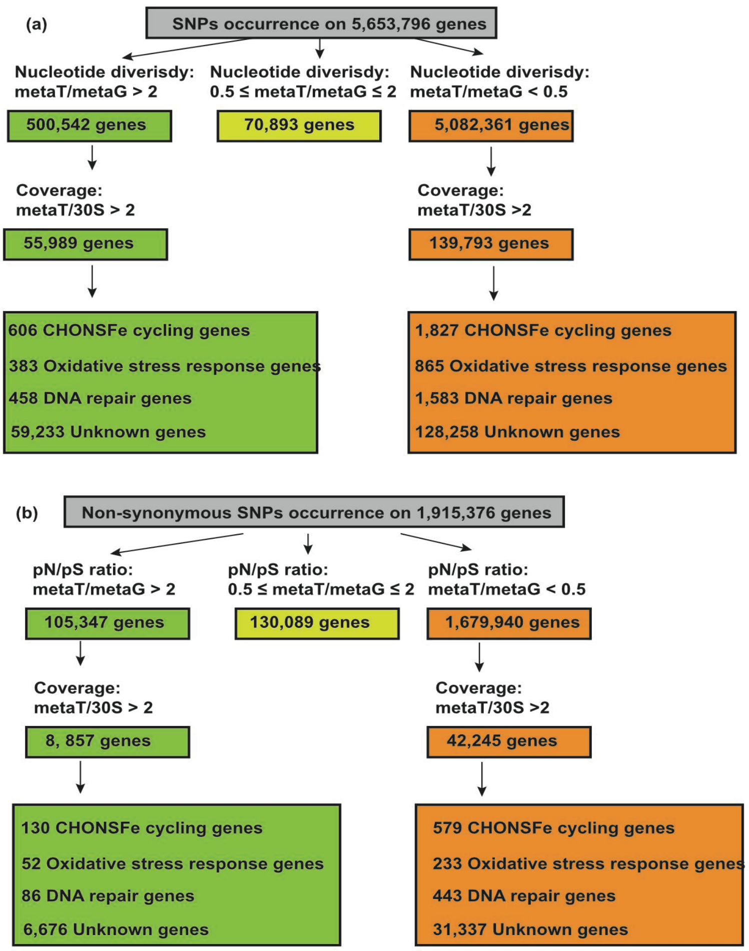

**Figure S19 | Distribution of in species-cluster rMAGs exhibiting signals of positive selection in metagenomic and metatranscriptomic datasets.** Venn diagram (top) shows the number of species-cluster representative MAGs (species-cluster rMAGs) with pN/pS ratios >1 in metatranscriptomes (MetaT), metagenomes (MetaG), or both. Pie charts (bottom) display the stage-wise distribution of the members of these species clusters with pN/pS ratios >1 across background (BG), mining (MT), and post-mining (PM) stages of neutral-pH ISL.

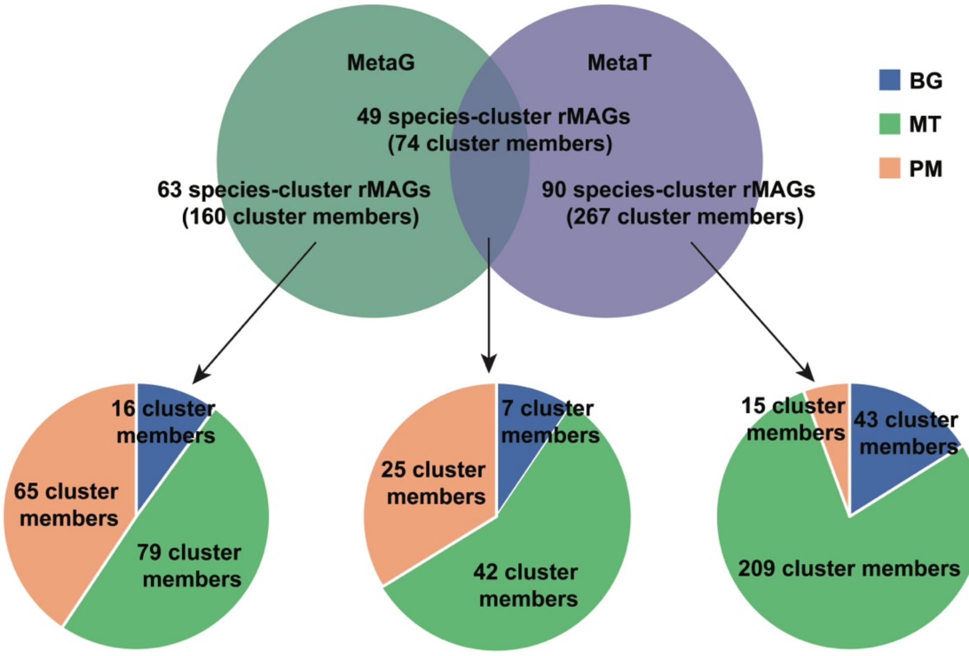

**Figure S20 | Proportions of rRNA/tRNA and retained non-rRNA reads after SortMeRNA filtering across mining phases.** Boxplots show the percentages of rRNA/tRNA reads (green) removed by SortMeRNA and the retained non-rRNA reads (blue) in metatranscriptomic datasets from background (BG), mining-impacted (MT), and post-mining (PM) aquifers.

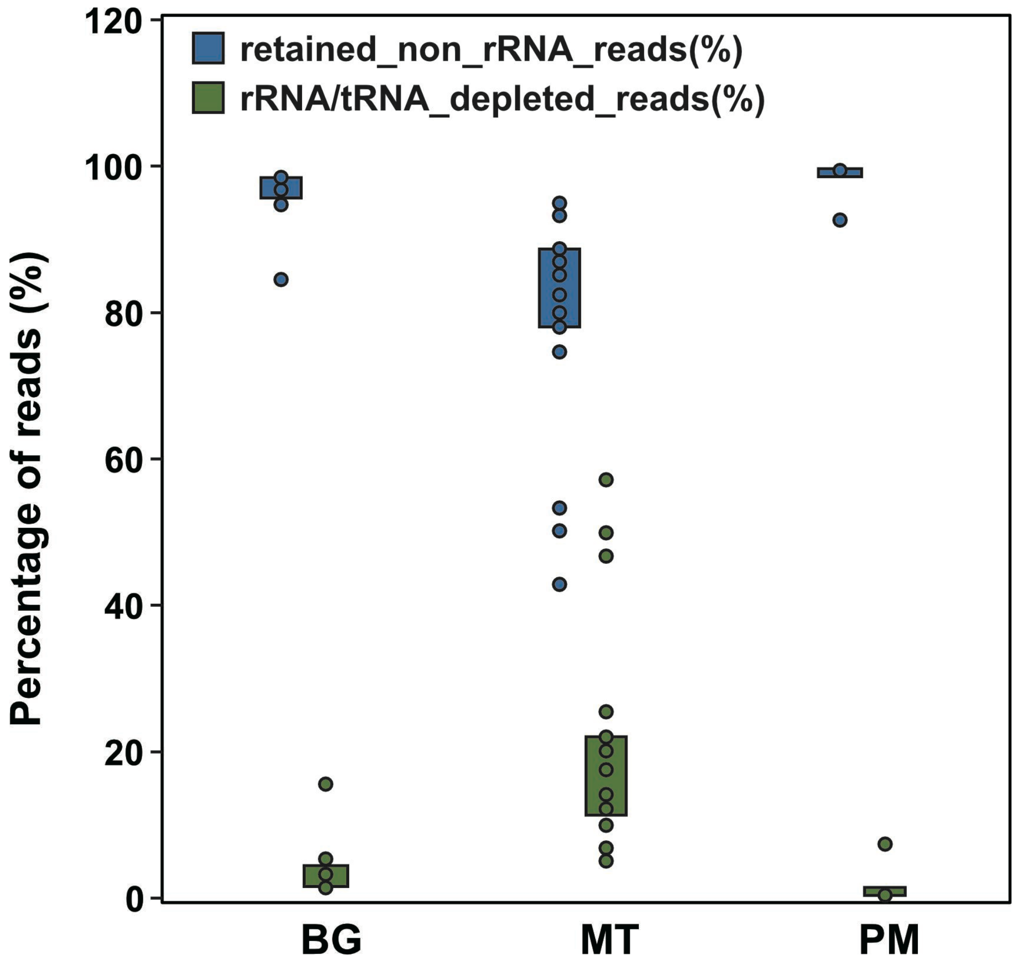
